# Reversible control of kinase signaling through chemical-induced dephosphorylation

**DOI:** 10.1101/2024.06.27.600945

**Authors:** Ying Sun, Rihong Zhou, Jin Hu, Shan Feng, Qi Hu

## Abstract

The coordination between kinases and phosphatases is crucial for regulating the phosphorylation levels of essential signaling molecules. Methods enabling precise control of kinase activities are valuable for understanding the kinase functions and for developing targeted therapies. Here, we use the abscisic acid (ABA)-induced proximity system to reversibly control kinase signaling by recruiting phosphatases. Using this method, we found that the oncogenic tyrosine kinase BCR::ABL1 can be inhibited by recruiting various cytoplasmic phosphatases. We also discovered that the oncogenic serine/threonine kinase BRAF(V600E), which has been reported to bypass phosphorylation regulation, can be positively regulated by protein phosphatase 1 (PP1) and negatively regulated by PP5. Additionally, we observed that the dual-specificity kinase MEK1 can be inhibited by recruiting PP5. This suggests that bifunctional molecules capable of recruiting PP5 to MEK or RAF kinases could be promising anticancer drug candidates. Thus, the ABA-induced dephosphorylation method enables rapid screening of phosphatases to precisely control kinase signaling.

## Introduction

Kinases and phosphatases regulate growth, differentiation and other essential cellular processes^1^. Protein kinases usually function as signal transducers by receiving activating signals from upstream ligands or regulators and catalyzing the phosphorylation of downstream substrate proteins; in contrast, phosphatases counteract the functions of kinases by catalyzing the dephosphorylation reaction. In human, 518 protein kinases and 189 protein phosphatases have been identified^2,3^. Dysregulation of kinases and phosphatases contributes to the development of diseases such as cancers^4^, neurodegenerative diseases^5^, and metabolic diseases^6^. How the protein kinases and phosphatases coordinate to maintain the phosphorylation of substrate proteins at proper levels is always a focus of kinase/phosphatase related studies.

In addition to sharing the same substrates, protein kinases and phosphatases also regulate the enzymatic activities of each other. In particular, the activities of protein kinases are tightly controlled by phosphorylation and dephosphorylation. For example, the oncogenic kinase BCR-ABL1, which is encoded by the fusion gene *BCR-ABL1* that is found in most chronic myelogenous leukemia (CML) patients, requires phosphorylation of its activation loop to be activated^7^. Following the recommendations from the HUGO Gene Nomenclature Committee (HGNC) to describe gene fusions^8^, we use the name BCR::ABL1 instead of BCR-ABL1 in this study. The RAF kinases, including CRAF, ARAF and BRAF, represent kinases that are regulated by more complicated phosphorylation and dephosphorylation processes^9^. Phosphorylation of BRAF at S365 inhibited the kinase activity, while phosphorylation at S729 promotes the activation of BRAF^9^.

Manipulating the phosphorylation and dephosphorylation of protein kinases or their substrates is a promising strategy to regulate the kinase signaling pathways. Lim et al. showed that the catalytic domain of a tyrosine phosphatase SHP1 with its N terminus fused to the ABL binding domain (ABD) of RIN1 could dephosphorylate BCR::ABL1 to inhibit its oncogenic activity^10^. Similarly, Simpson et al. reported that phosphatases fused to an anti-GFP nanobody could bind to GFP-fused target proteins and dephosphorylate the target proteins^11^.

Inspired by proteolysis targeting chimeras (PROTACs)^12^, which consist of a binder or inhibitor of the target protein, a binder or activator of an E3 ubiquitin ligase, and a linker covalently linked the two binders, Yamazoe et al. developed phosphatase recruiting chimeras (PhoRCs) by covalently linking a binder of the target protein with a binder or activator of a phosphatase, and demonstrated that PHORCs effectively inhibited AKT by recruiting protein phosphatase 1 (PP1) to dephosphorylate AKT^13^. The same strategy has been used to design heterobifunctional molecules that dephosphorylated Tau by recruiting protein phosphatase 2A (PP2A)^14^, dephosphorylated FOXO3a by PP2A^15^, and that dephosphorylated ASK1 by protein phosphatase 5 (PP5)^16^, highlighting the therapeutic potential of the PHORC strategy. But given the large number of protein kinases and phosphatases and their complicated regulatory mechanisms, a key question for the PHORC strategy is which phosphatase should be chosen to dephosphorylate a given target protein. In this study, using the chemically induced proximity (CIP) strategy we develop a method to quickly screen phosphatases to precisely control kinase signaling. The basic idea of the CIP strategy is to use a chemical to induce the proximity of two or more molecules, especially biomacromolecules^17^. Several CIP systems have been identified^17^. For example, the rapamycin-induced dimerization system has been widely used in biological studies, such as to regulate gene expression^18^, and protein translocation^19,20^. However, the interference of rapamycin with mTOR and FKBP12 limits its application. In contrast, the plant hormone abscisic acid (ABA) induced proximity system, in which ABA induces the binding of plant PYR1/PYL/RCAR proteins (PYLs) to a plant phosphatase ABI1, has the advantage of not interacting with proteins in mammalian cells^21^. ABA has good stability and bioavailability, and lower toxicity in mammalian cells^21^. A point mutation — D143A was introduced into ABI1 to disrupt the phosphatase activity of ABI1^21^. We fused the D143A mutant of ABI1 to the N or C termini of kinases, fused PYL2 to the N or C termini of phosphatases, and then used ABA to induce the proximity of the kinases and phosphatases. We demonstrated that the ABA induced proximity system can be used to reversibly control both tyrosine kinases (BCR::ABL1 as an example) and serine/threonine kinases (BRAF-MEK-ERK pathway as an example). Using this method, we identified phosphatases that can be recruited to regulate BCR::ABL1, BRAF(V600E) and MEK1.

## Results

### The ABA induced proximity system can reversibly control BCR::ABL1 signaling

To test whether the ABA induced proximity system can be used to control tyrosine kinases, we chose BCR::ABL1 as an example. ABL contains a Src homology 3 (SH3) domain, a Src homology 2 (SH2) domain and a catalytic domain; in its unphosphorylated state, ABL has its catalytic domain inhibited by the SH2 and SH3 domains; fusion of the BCR (breakpoint cluster region) protein to the N-terminus of ABL leads to the dimerization and phosphorylation of BCR::ABL1, resulting in the release of the SH2 and SH3 domains from the catalytic domain and constitutive activation of ABL. As for the phosphatase, we chose the tyrosine phosphatase SHP1 that has been reported to inhibit the function of BCR::ABL1. By fusing a His-tagged PYL2 (residues 11 to 188) to the N terminus of SHP1 and by fusing a FLAG-tagged ABI (ABI1, residues 126 to 422, with the D143A mutation) to the N terminus of BCR::ABL1, we created two chimeric proteins His-PYL2-SHP1 and FLAG-ABI-BCR::ABL1 (Figure 1A).

**Figure 1.**
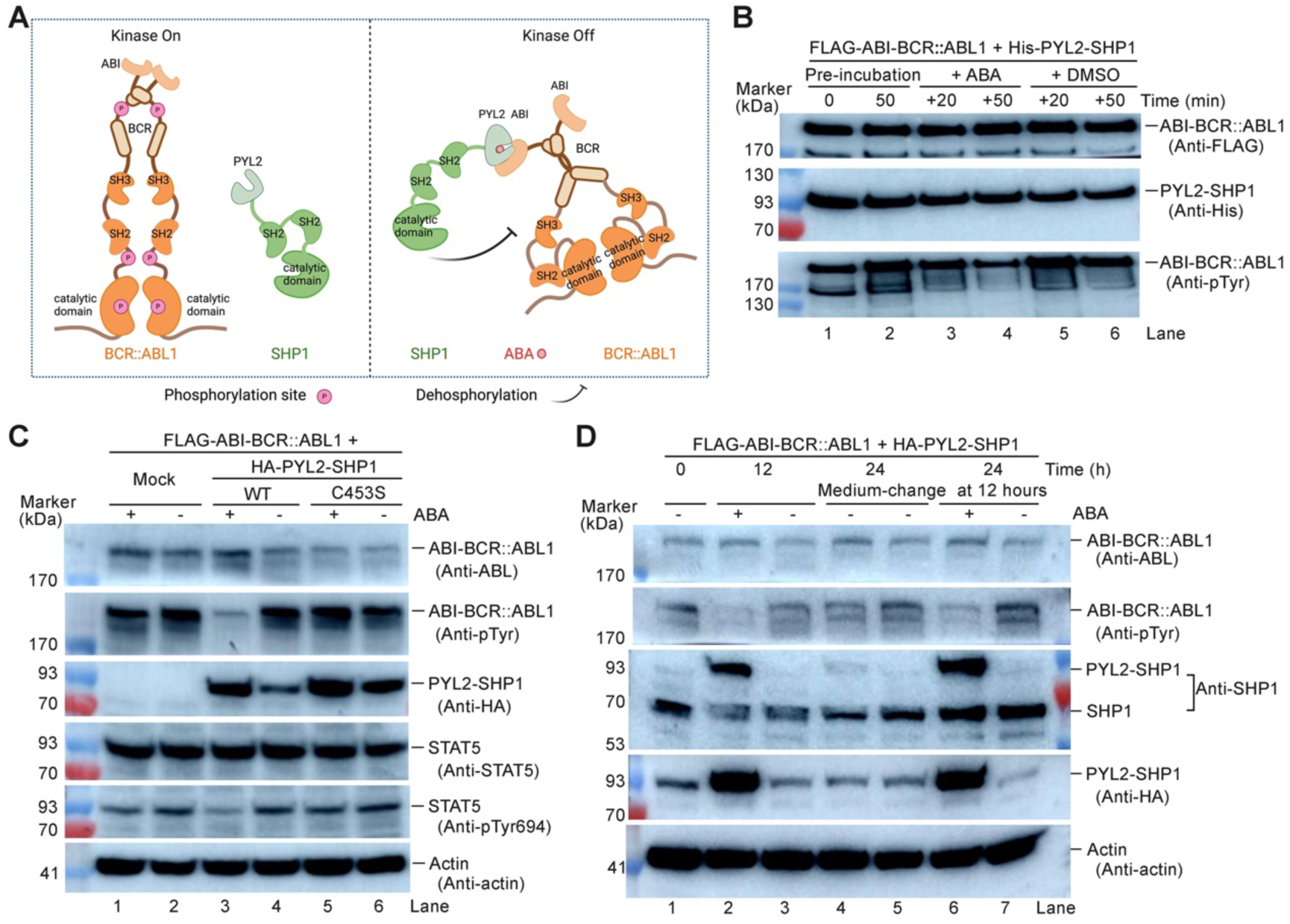
Recruitment of SHP1 to BCR::ABL1 dephosphorylated BCR::ABL1 and inhibited its activity to phosphorylate STAT5. (A) Illustration of the ABA induced proximity system. ABI (residues 126-422, D143A) and PYL2 (residues 11-188) are fused to the N termini of BCR::ABL1 and SHP1, respectively. ABA induces the dimerization of PYL2 with ABI to bring SHP1 close to BCR::ABL1. (B) The purified FLAG-tagged ABI-BCR::ABL1 could be phosphorylated by itself and dephosphorylated by recruiting His-tagged PYL2-SHP1 to it. (C) In Ba/F3 cells, recruitment of HA-tagged PYL2-SHP1 decreased the phosphorylation level of FLAG-tagged ABI-BCR::ABL1 and inhibited the phosphorylation of STAT5 at Y694. (D) The decreased phosphorylation level of FLAG-tagged ABI-BCR::ABL1 by recruiting HA-tagged PYL2-SHP1 in Ba/F3 cells was recovered after ABA was withdraw.

We overexpressed the two chimeric proteins and purified them to homogeneity. First, we confirmed that ABA can induce the interaction between His-PYL2-SHP1 and FLAG-ABI-BCR::ABL1 using a gel filtration assay (Figure S1). Next, we assessed the impact of His-PYL2-SHP1 on the autophosphorylation level of FLAG-ABI-BCR::ABL1. After incubation of the purified His-PYL2-SHP1 with FLAG-ABI-BCR::ABL1 in a kinase reaction buffer for 50 minutes, an increase of the tyrosine phosphorylation (pTyr) level of FLAG-ABI-BCR::ABL1 was observed (Figure 1B, lanes 1 vs. 2). Then half of the reaction was incubated with ABA while the other half was incubated with DMSO for additional 20 or 50 minutes. Upon addition of ABA but not DMSO, the pTyr level of BCR::ABL1 sharply decreased (Figure 1B, lanes 3 to 6), suggesting that recruitment of SHP1 to BCR::ABL1 by the ABA induced proximity system can promote the dephosphorylation of BCR::ABL1.

Next, we tested whether the ABA induced proximity system can control BCR::ABL1 signaling in Ba/F3 cells. Ba/F3 cells are a murine pro-B cell line frequently used in the study of oncogenic kinases due to their dependence on interleukin-3 (IL-3) for survival, which can be overridden by oncogenic mutations in the kinases^22^. We stably expressed FLAG-tagged ABI-BCR::ABL1 alone or together with HA-tagged PYL2-SHP1 in Ba/F3 cells. When expressed together with wild-type PYL2-SHP1 but not expressed alone or expressed together with PYL2-SHP1 carrying a catalytic-dead mutation C453S in SHP1, ABI-BCR::ABL1 had its pTyr level decreased upon treatment with ABA (Figure 1C). The pTyr level of STAT5, which is an important substrate of BCR::ABL1, was also decreased by the recruitment of SHP1 to BCR::ABL1 (Figure 1C, lanes 3 vs. 4).

In addition to STAT5, BCR::ABL1 has also been reported to be able to activate the PI3K/AKT and MAPK/ERK pathways^23^, and catalyze the phosphorylation of the SH2/SH3 domain-containing protein CrkL^24^. Therefore, we also checked changes in the phosphorylation levels of AKT, ERK and CrkL in Ba/F3 cells upon treatment with ABA or imatinib (IMA). IMA is a small molecule inhibitor of BCR::ABL1 and has been approved for the treatment of CML^25^. In contrast to the obvious decrease in the phosphorylation level of STAT5, the phosphorylation levels of AKT, ERK, and CrkL showed little or moderate change upon treatment with ABA or IMA for 12 h (Figure S2).

The decrease of pTyr level of BCR::ABL1 is reversible. In Ba/F3 cells stably expressing both ABI-BCR::ABL1 and PYL2-SHP1, after 12-hour treatment with ABA, the pTyr level of BCR::ABL1 was decreased (Figure 1D, lanes 2 vs. 3). Then for half of the cells, the medium was changed to fresh medium not containing ABA, and after additional 12 hours, the pTyr level of BCR::ABL1 was recovered (Figure 1D, lane 4); in the other half of the cells of which the medium was changed to fresh medium containing ABA, the pTyr level of BCR::ABL1 was still very low after additional 12 hours (Figure 1D, lane 6). These results suggest that the ABA induced proximity system can reversibly control BCR::ABL1 signaling in Ba/F3 cells.

### ABA can increase the protein level of PYL2 fusion proteins

Unexpectedly, we found that the protein level of PYL2-SHP1 was increased upon treatment with ABA (Figure 1C, lanes 3 and 5; Figure 1D, lanes 2 and 6). A similar result was also observed when PYL2-SHP1 was co-expressed with FLAG-tagged ABI (Figure S3A) or with BCR::ABL1 (Figure S3B). But when the interaction between PYL2 and ABA was disrupted by introducing a K64A mutation into PYL2, adding ABA could no longer increase the protein level of PYL2-SHP1 (Figure S3C). Even if there was no SHP1 fused to PYL2, adding ABA could still increase the protein level of PYL2 (Figure S3D). We conclude that the ABA induced upregulation of PYL2-SHP1 was not because of the function of BCR::ABL1, SHP1, or the interaction between ABI and PYL2, but because of the binding of ABA to PYL2. This unexpected stabilizing effect of ABA binding further adds to the inducibility of this CIP system.

To test whether the degradation of PYL2 occurs through the ubiquitin-proteasome pathway, we treated the Ba/F3 cells with MG132, a proteasome inhibitor^26^. Unexpectedly, MG132 treatment decreased the protein levels of both FLAG-tagged ABI-BCR::ABL1 and HA-tagged PYL2-SHP1 (Figure S3E). A previous study also found that the protein level of BCR::ABL1 decreased upon MG132 treatment, but the reason is unclear^27^.

### Screening of phosphatases to regulate BCR::ABL1 phosphorylation

In addition to SHP1, we speculate whether other phosphatases can also be used to regulate BCR::ABL1 phosphorylation. The human protein phosphatases, which can be divided into serine/threonine phosphatases (PSPs), tyrosine phosphatases (PTPs), and dual-specificity phosphatases (DUSPs), have various subcellular locations. We selected several representative phosphatases to test using the ABA induced proximity system (Figure 2A).

**Figure 2.**
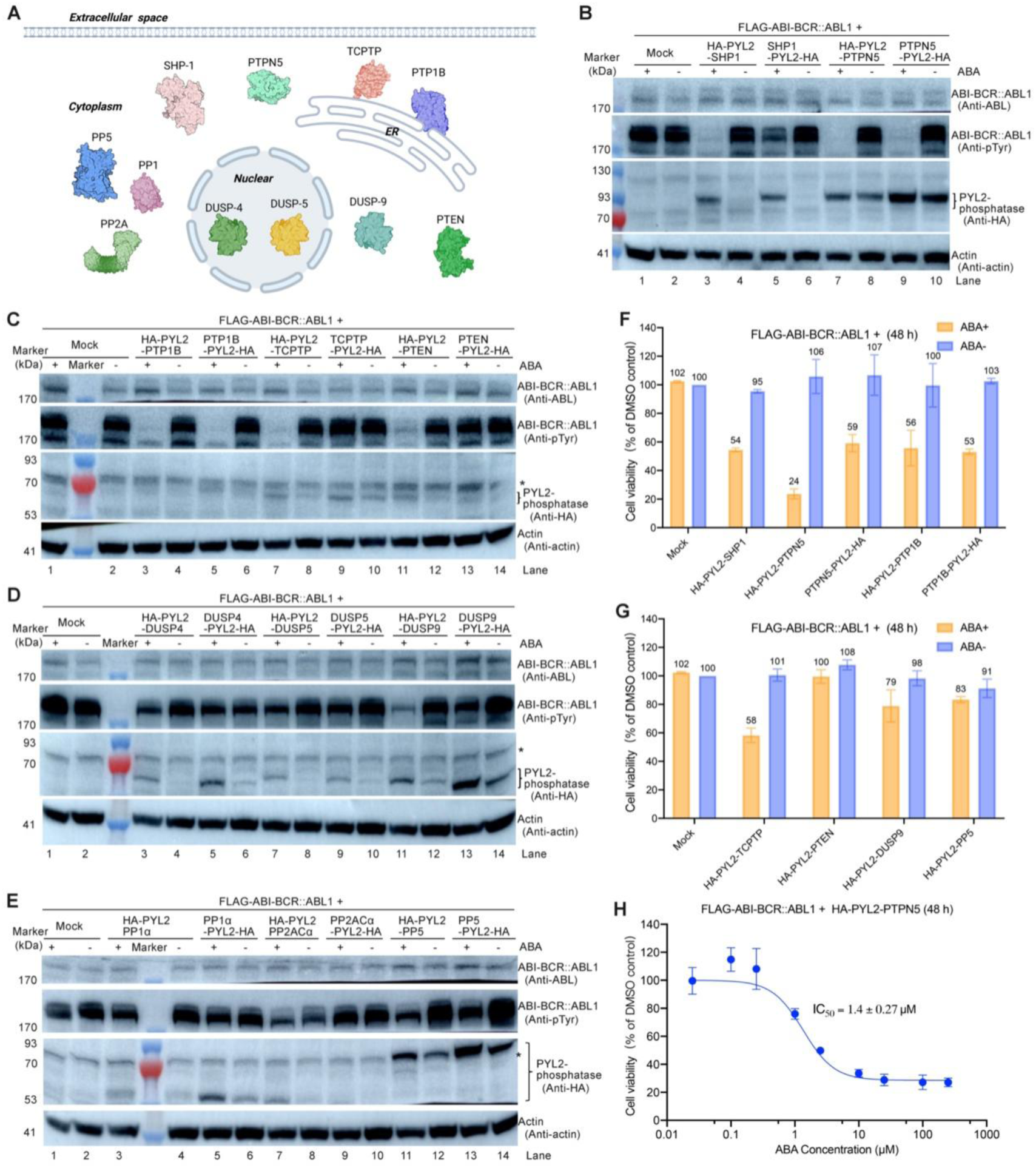
Identification of phosphatases that can dephosphorylate BCR::ABL1 when being recruited to BCR::ABL1. (A) Illustration of the subcellular localization of representative phosphatases in human cells. (B-E) Screening of phosphatases that can be recruited to BCR::ABL1 to regulate BCR::ABL1 phosphorylation in Ba/F3 cells using the ABA induced proximity system. The asterisk (*) indicates a non-specific band detected by the anti-HA antibody. (F and G) Inhibition of the viability of Ba/F3 cells expressing FLAG-tagged ABI-BCR::ABL1 by recruitment of different phosphatases to ABI-BCR::ABL1 using ABA. The cell viability of each group was normalized to that of the ABA-untreated mock group. The values represent the mean ± SD of two independent measurements with technical duplicates. (H) ABA inhibited the viability of Ba/F3 cells expressing FLAG-tagged ABI-BCR::ABL1 and HA-tagged PYL2-PTPN5, with an IC_50_ of 1.4 µM. The values represent the mean ± SD of two independent measurements with technical duplicates.

The first type of phosphatases we tested are PTPs. SHP1 and a cytosolic variant of PTPN5 (also called STEP, STriatal-Enriched protein tyrosine Phosphatase) are cytosolic PTPs; PTP1B and TCPTP are also PTPs, but are localized to the endoplasmic reticulum (ER). Previous studies have showed that PTP1B can dephosphorylate BCR::ABL1 at Tyr1086 to stabilize BCR::ABL1^28^. We fused HA-tagged PYL2 to the N or C termini of these phosphatases, and stably expressed them together with FLAG-tagged ABI-BCR::ABL1 in Ba/F3 cells. After 12-hour treatment with ABA, a significant decrease of the pTyr level of FLAG-ABI-BCR::ABL1 was observed in Ba/F3 cells expressing HA-PYL2-SHP1, HA-PYL2-PTPN5, PTPN5-PYL2-HA, HA-PYL2-PTP1B, PTP1B-PYL2-HA, or HA-PYL2-TCPTP (Figures 2B and 2C).

Next, we tested DUSPs, which can dephosphorylate a variety of substrates including protein tyrosine, threonine, and serine residues, and phosphoinositides^29^. We chose PTEN and DUSP9 as representatives of cytosolic DUSPs, and DUSP4 and DUSP5 as representatives of DUSPs in the nucleus. PTEN prefers phosphatidylinositol (3,4,5)-trisphosphate (PIP_3_) as its substrate, but can also dephosphorylate proteins^30^. DUSP9 has been reported to dephosphorylate mitogen-activated protein kinases (MAPKs)^31^. DUSP4 and DUSP5 are known to dephosphorylate extracellular signal-regulated kinases (ERKs) in nucleus^32^. We found that only PTEN and DUSP9 with HA-tagged PYL2 fused to their N termini decreased the pTyr level of ABI-BCR::ABL1 upon treatment with ABA (Figures 2C and 2D).

Lastly, we tested three cytosolic PSPs, including PP1, PP2A, and PP5. A previous study showed that recruiting PP1 to EGFR reduced the phosphorylation level of EGFR at Y1068, suggesting that PSPs could be promiscuous in dephosphorylating tyrosine residues^13^. We found that only recruitment of PP5 induced the dephosphorylation of BCR::ABL1 (Figure 2E).

By using catalytic-dead mutants of PTPN5, PTP1B, TCPTP, and PTEN as controls, we demonstrated that the effects of recruiting these phosphatases to BCR::ABL1 on the pTyr level of BCR::ABL1 depend on the catalytic activities of these phosphatases (Figure S4). For PTPN5, recruitment of its catalytic-dead mutant to BCR::ABL1 also decreased the pTyr level of BCR::ABL1, though the decrease was not as significant as that caused by recruitment of active PTPN5, indicating that non-catalytic effect of PTPN5 also contributes to its inhibitory activity against BCR::ABL1 phosphorylation.

We next tested the effect of ABA on the cell viability of Ba/F3 cells stably expressing both FLAG-tagged ABI-BCR::ABL1 and HA-tagged PYL2-phosphatase fusion proteins. ABA had the highest activity in Ba/F3 cells expressing HA-PYL2-PTPN5 (Figures 2F and 2G), with an IC_50_ value of 1.4 µM (Figure 2H).

The above results show that PTPs and DUSPs in cytoplasm can dephosphorylate BCR::ABL1 when they are recruited to BCR::ABL1 through the ABA induced proximity system. Only when PYL2 was fused to the N termini but not C termini of SHP1, TCPTP, PTEN, and DUSP9, recruitment of these phosphatases to BCR::ABL1 dephosphorylated BCR::ABL1, indicating that the position and orientation of phosphatases relative to kinases should be taken into consideration when the ABA induced proximity system is used to control kinase signaling pathways. Regarding the phosphatases that fail to dephosphorylate BCR::ABL1, we cannot rule out the possibility that this is due to an improper distance or orientation between these phosphatases and BCR::ABL1 when brought together. Alternatively, it may be because of direct interference with the phosphatase activities caused by the fused HA-tagged PYL2.

### Synergism between ABA and the BCR::ABL1 inhibitor imatinib (IMA)

In line with the inhibitory effect of ABA on the proliferation of Ba/F3 cells, ABA reduced the pTyr level of BCR::ABL1 in a dose-dependent manner (Figure 3A). We compared the activities of ABA with IMA. IMA inhibited ABI-BCR::ABL1 phosphorylation regardless of the presence of PYL2-PTPN5. In contrast, the activity of ABA was dependent on the presence of PYL2-PTPN5 (Figure 3B).

**Figure 3.**
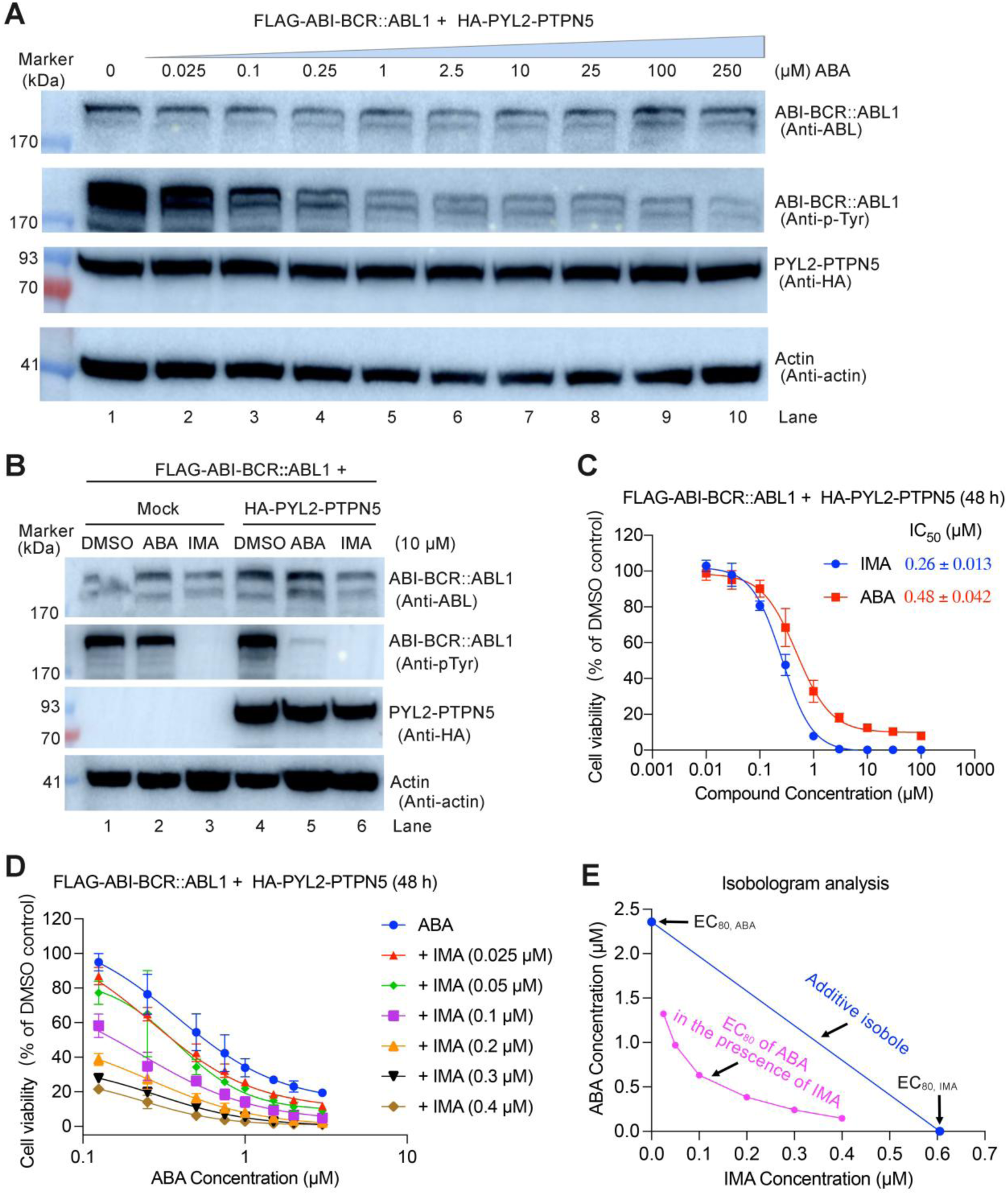
Regulation of the pTyr level of BCR::ABL1 in Ba/F3 cells using ABA and imatinib (IMA). (A) Dose-dependent effect of ABA on the pTyr level of BCR::ABL1 in Ba/F3 cells expressing FLAG-tagged ABI-BCR::ABL1 and HA-tagged PYL2-PTPN5. (B) Effects of IMA and ABA (10 µM) on the pTyr level of BCR::ABL1 in Ba/F3 cells expressing FLAG-tagged ABI-BCR::ABL1 alone or together with HA-tagged PYL2-PTPN5. (C-D) Inhibition of the proliferation of Ba/F3 cells expressing FLAG-tagged ABI-BCR::ABL1 and HA-tagged PYL2-PTPN5 by IMA or ABA alone (C), or combinations of different concentrations of ABA and IMA (D). The data represent the mean ± SD of three independent measurements with technical duplicates. (E) Isobologram analysis reveals synergism between ABA-induced dephosphorylation and inhibition of the kinase activity of BCR::ABL1 by IMA. The line called additive isobole was plotted using the equation “x/EC_80,IMA_ + y/EC_80,ABA_ = 1”, where “x” and “y” represent the concentrations of IMA and ABA, respectively, used in the combination treatment, while “EC_80,IMA_” and “EC_80,ABA_” represent the EC_80_ values of IMA and ABA, respectively, when they were used alone (panel C). The EC_80_ values of ABA in the presence of different concentrations of IMA (panel D) were calculated and used to plot the EC_80_ curve.

To test whether ABA-induced dephosphorylation has a synergistic effect with direct inhibition of the kinase activity of BCR::ABL1 by IMA, we treated the Ba/F3 cells expressing FLAG-tagged ABI-BCR::ABL and HA-tagged PYL2-PTPN5 with either ABA or IMA alone (Figure 3C), as well as with combinations of different concentrations of ABA and IMA (Figure 3D). We then conducted isobologram analysis^33^, to evaluate the presence of synergism (Figure 3E). The EC_80_ curve of ABA in the presence of different concentrations of IMA is below the additive isobole, indicating synergism between ABA and IMA.

### BRAF(V600E) is sensitive to ABA induced dephosphorylation

Having found that recruitment of phosphatases to BCR::ABL1 dephosphorylated BCR::ABL1, we next tested using the ABA induced proximity system to control serine/threonine kinases. We focused on the RAF-MEK-ERK kinase signaling pathway that plays a pivotal role in the regulation of cell growth, differentiation, and survival^34^. Aberrant activation of this pathway contributes to the initiation and progression of many cancers^35^. Among the RAF kinases, BRAF carries the most prevalent oncogenic RAF mutation — BRAF V600E, which has been found in approximately 8% of human tumors, particularly in melanoma, colorectal, thyroid cancers^36,37^. We tried to regulate the kinase activity of the BRAF(V600E) mutant using three serine/threonine phosphatases, including PP1α, PP2ACα and PP5. PP1α and PP2ACα are the catalytic subunits of PP1 and PP2A, respectively. Previous studies have shown that the PP1 and PP2A can positively regulate wild-type RAF kinases^38,39^, while PP5 plays a negative role^40^.

We fused the FLAG-tagged ABI to the C terminus of BRAF(V600E) and fused the HA-tagged PYL2 to the N or C termini of the phosphatases, then stably expressed them in Ba/F3 cells to make the cells dependent on the RAF-MEK-ERK pathway after IL-3 withdrawal. ABA-induced recruitment of HA-tagged PYL2-PP1α and PYL2-PP2ACα to FLAG-tagged BRAF(V600E)-ABI was confirmed by purifying the kinase-phosphatase complexes using immobilized anti-FLAG antibody and subsequently conducting mass spectrometry (MS) analysis (Table S1). In addition to PP1α and PP2ACα, several other subunits of PP1 and PP2A were also co-purified with FLAG-tagged BRAF(V600E)-ABI in the presence of ABA (Table S1).

PP1α, PP2ACα and PP5 showed distinct effects on the BRAF(V600E)-MEK-ERK signaling pathway (Figure 4A). Recruitment of PP1α did not have an apparent effect on the phosphorylation of S729 and S446 of BRAF, nor on the phosphorylation of S218/S222 of MEK, but increased the phosphorylation level of T202/Y204 of ERK (Figure 4A, lanes 3-6). Recruitment of PP2ACα had little effect on the phosphorylation level of BRAF, MEK or ERK (Figure 4A, lanes 7-10). In contrast, recruitment of PP5 dephosphorylated BRAF(V600E) at S729 and S446, inhibiting the phosphorylation of MEK S218/S222 and ERK T202/Y204 (Figure 4A, lanes 11-14). We further confirmed that the distinct effects of PP1α and PP5 on ERK phosphorylation were attributed to the phosphatase activities of PP1α and PP5 by introducing catalytic-dead mutations H248K and H304A into PP1α and PP5, respectively (Figures 4B and 4C). Recruitment of PP5 to BRAF(V600E) significantly inhibited the viability of the Ba/F3 cells (Figure 4D). Similar results were observed when we fused the FLAG-tagged ABI to the N terminus of BRAF(V600E) (Figure S5). These results demonstrate that PP1α and PP5 have opposite effects on the RAF-MEK-ERK signaling pathway when they are recruited to BRAF(V600E); in addition, recruitment of PP5 to BRAF(V600E) is a feasible strategy to inhibit BRAF(V600E) activity (Figure 4E).

**Figure 4.**
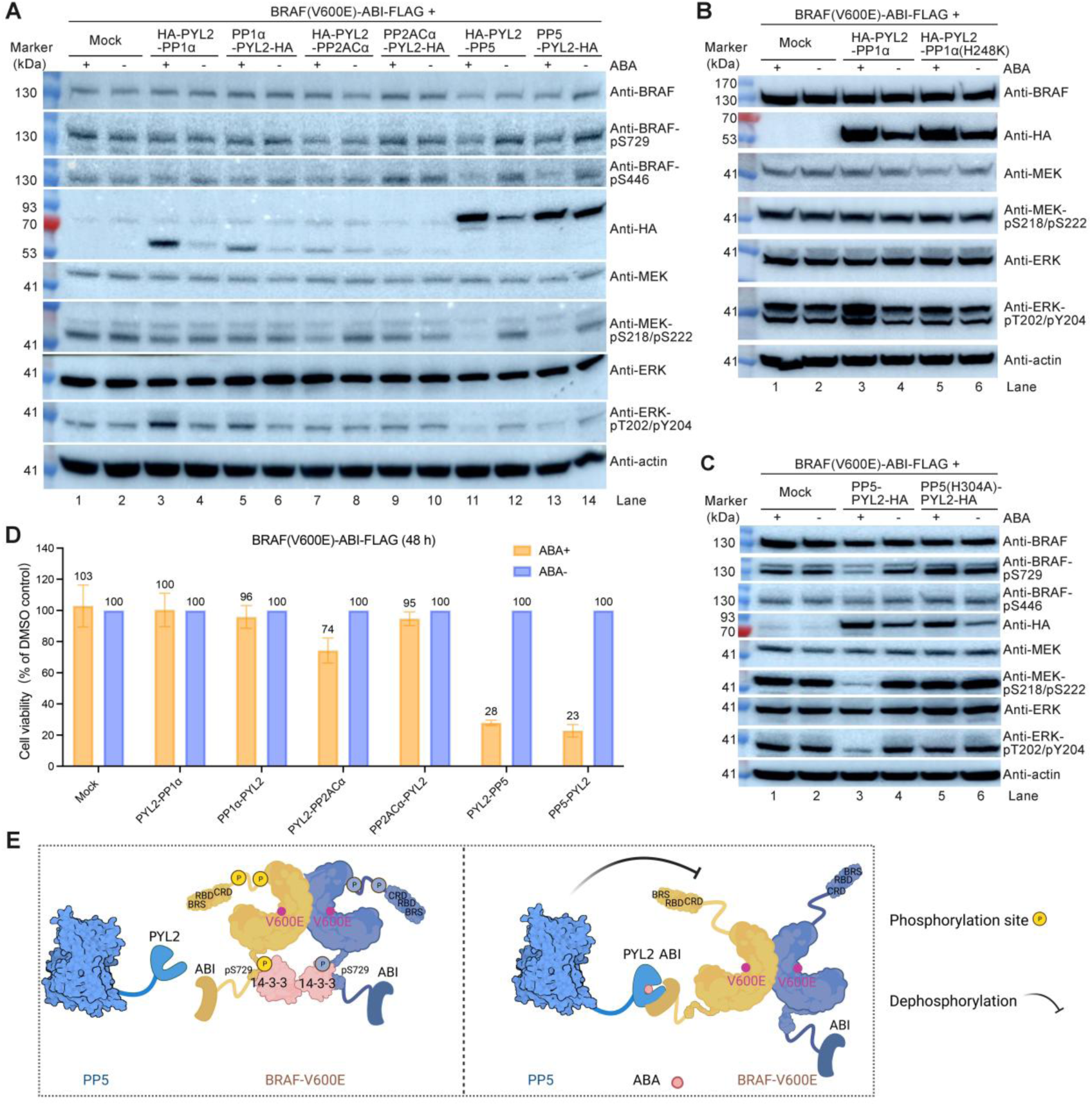
Regulation of BRAF(V600E) activity by recruiting protein serine/threonine phosphatases. (A) Recruitment of PP1α, PP2ACα and PP5 to BRAF(V600E) using the ABA induced proximity system in Ba/F3 cells showed distinct effects on the BRAF(V600E)-MEK-ERK signaling pathway. (B) Recruitment of wild-type PP1α but not the catalytic-dead mutant PP1α(H248K) to BRAF(V600E) in Ba/F3 cells increased the phosphorylation level of ERK. (C) Recruitment of wild-type PP5 but not the catalytic-dead mutant PP5(H304A) to BRAF(V600E) in Ba/F3 cells blocked the activation of MEK and ERK. (D) ABA inhibited the viability of Ba/F3 cells expressing FLAG-tagged BRAF(V600E)-ABI and HA-tagged PP5-PYL2, with an IC_50_ of 3.0 µM. The values represent the mean ± SD of two independent measurements with technical duplicates. (E) Illustration of the regulation of BRAF(V600E) by PP5 using the ABA induced proximity system.

The different effects of PP1α and PP5 on BRAF(V600E) signaling pathway are consistent with their effects on wild-type RAF signaling pathway reported by previous studies, in which it was found that PP1 dephosphorylated CRAF at S259 (corresponding to S365 in BRAF) to promote CRAF activation^38,39^, while PP5 dephosphorylated CRAF at S338 (corresponding to S446 in BRAF) to inhibit CRAF activity^40^. But it has been reported that the oncogenic mutation V600E allows BRAF to bypass the regulation by phosphorylation/dephosphorylation^41–45^. To understand the effects of PP1α and PP5 on BRAF(V600E), we harvested the Ba/F3 cells with and without ABA treatment (Figure S6A), enriched the FLAG-tagged BRAF(V600E) using anti-FLAG beads and quantified its phosphorylation using mass spectrometry (MS). In Ba/F3 cells expressing both FLAG-ABI-BRAF(V600E) and HA-PYL2-PP5, addition of ABA induced dephosphorylation of BRAF(V600E) at S319/S325, S365, S399/T401, and S729; dephosphorylation of these residues was also observed in Ba/F3 cells expressing both BRAF(V600E)-ABI-FLAG and PP5-PYL2-HA, but to a less extent (Figures S6B to S6G). In Ba/F3 cells expressing both FLAG-ABI-BRAF(V600E) and HA-PYL2-PP1α, adding ABA decreased the phosphorylation at S365, S399/T401, and S729, but the decrease was not as significant as that in the presence of HA-PYL2-PP5 and ABA (Figures S6D, S6E, and S6G).

### Drug-resistant variants of BRAF(V600E) can be inhibited by recruiting PP5

Three BRAF inhibitors, including vemurafenib (VEM), dabrafenib, and encorafenib, have received approval for the treatment of cancers, such as melanomas and non-small cell lung cancers (NSCLC), that carry the BRAF V600E or V600K mutation^46^. Vemurafenib was approved for the treatment of BRAF-mutated melanomas in 2011 in the United States^47^, but soon drug-resistant variants of BRAF(V600E) were identified.^48^ Melanoma cell line SKMEL-239 carrying aberrant splicing variants of BRAF(V600E) lack the RAS binding domain showed strong resistance to VEM (Figure 5A)^48^. aberrant spliced BRAF(V600E) also exhibited resistance to dabrafenib and encorafenib^49,50^. We confirmed that in Ba/F3 cells co-expressing these spliced BRAF(V600E) and PP5 were resistant to VEM (Figures 5B and 5C). The IC_50_ value of VEM against the full-length BRAF(V600E) was 0.4 µM but increased to around 30 µM against the p61 and p55 variants (Figure 5B). In contrast, aberrant splicing of BRAF(V600E) did not confer resistance to ABA-induced dephosphorylation (Figures 5D and 5E). These results demonstrate the potential of the induced dephosphorylation strategy in overcoming drug resistance caused by alterations of the kinases.

**Figure 5.**
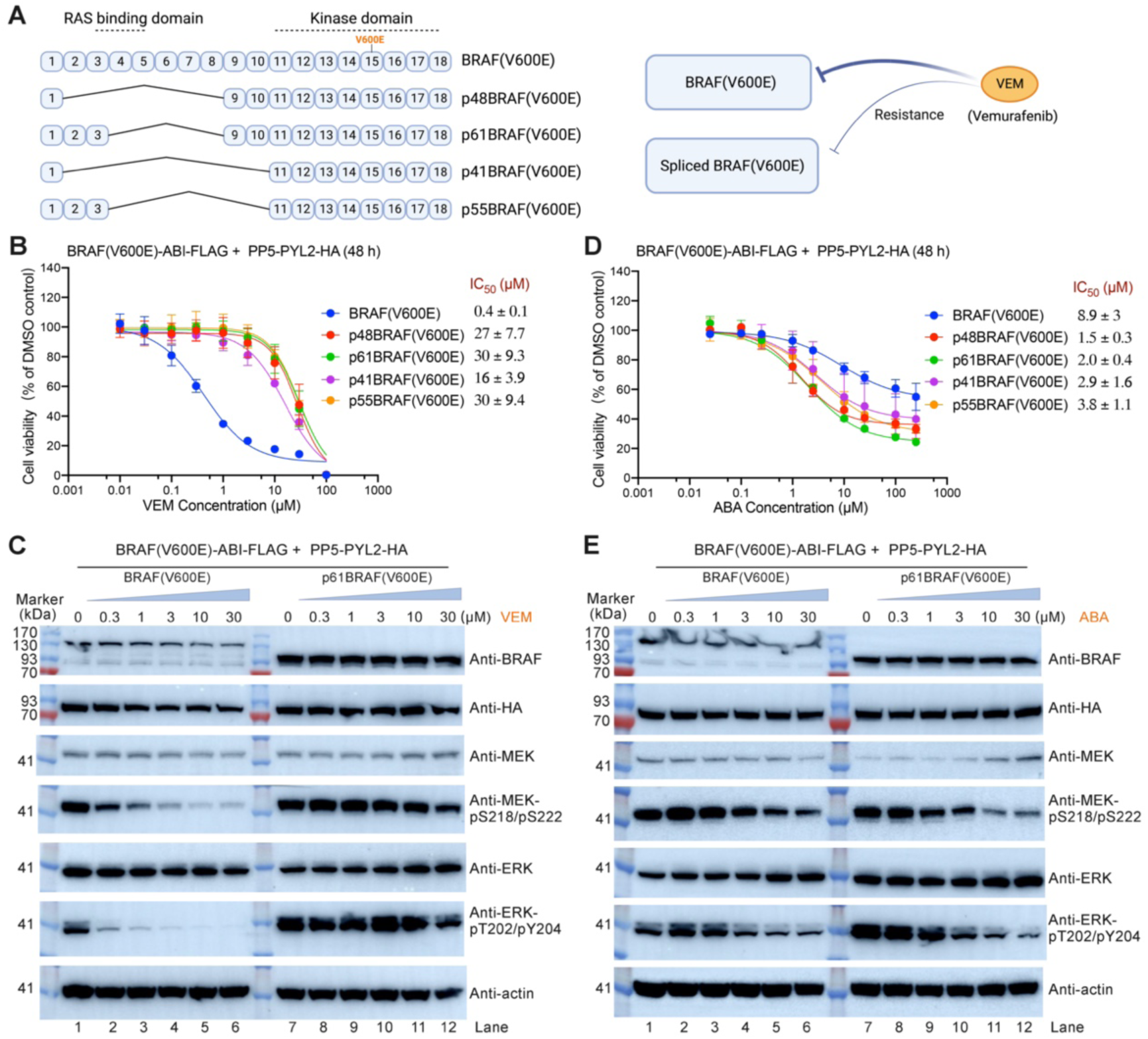
Splicing variants of BRAF(V600E) resistant to vemurafenib (VEM) are sensitive to ABA-induced dephosphorylation. (A) Illustration of the exon organization of the aberrant splicing variants of BRAF(V600E) that are resistant to vemurafenib (VEM). (B and D) Ba/F3 cells expressing full-length or aberrantly spliced BRAF(V600E)-ABL-FLAG together with PP5-PYL2-HA were treated with VEM (B) or ABA (D), and the cell viability was measured after 48 h. The data represents the mean ± SD of two independent measurements, each with technical duplicates. (C and E) Effects of VEM (C) and ABA (E) on the phosphorylation levels of MEK and ERK in Ba/F3 cells expressing BRAF(V600E)-ABI-FLAG or p61BRAF(V600E)-ABI-FLAG together with PP5-PYL2-HA.

### Recruitment of PP5 to MEK1 can inactivate MEK1

The dual-specificity kinases MEK1 and MEK2 are at the center of the RAF-MEK-ERK signaling pathway^51^. Human MEK1 shares a sequence identity of 80% with MEK2. The functions of MEK1 and MEK2 are redundant^52^. Therefore, we only focused on MEK1. We tested whether the ABA induced proximity system can regulate MEK1 activity. We fused FLAG-tagged ABI to the C terminus of MEK1, fused HA-tagged PYL2 to the N or C termini of PP1α, PP2ACα and PP5, then stably expressed them in Ba/F3 cells stably expressing BRAF(V600E). In the presence of ABA, HA-tagged PYL2-PP1α, PYL2-PP2ACα, and several other subunits of PP1 and PP2A were co-purified with FLAG-tagged MEK-ABI (Table S1), indicating that ABA can promote the kinase-phosphatase interactions. Upon treatment with ABA, only in cells expressing HA-PYL2-PP5 we observed the decrease of both MEK-ABI phosphorylation and ERK phosphorylation (Figure 6A). When the catalytic-dead mutation H304A was introduced into PP5, recruitment of HA-PYL2-PP5 could not decrease the phosphorylation level of either MEK or ERK (Figure 6B). These results demonstrate that MEK1 can be inactivated by recruiting PP5 to it, and the phosphatase activity of PP5 is required (Figure 6C).

**Figure 6.**
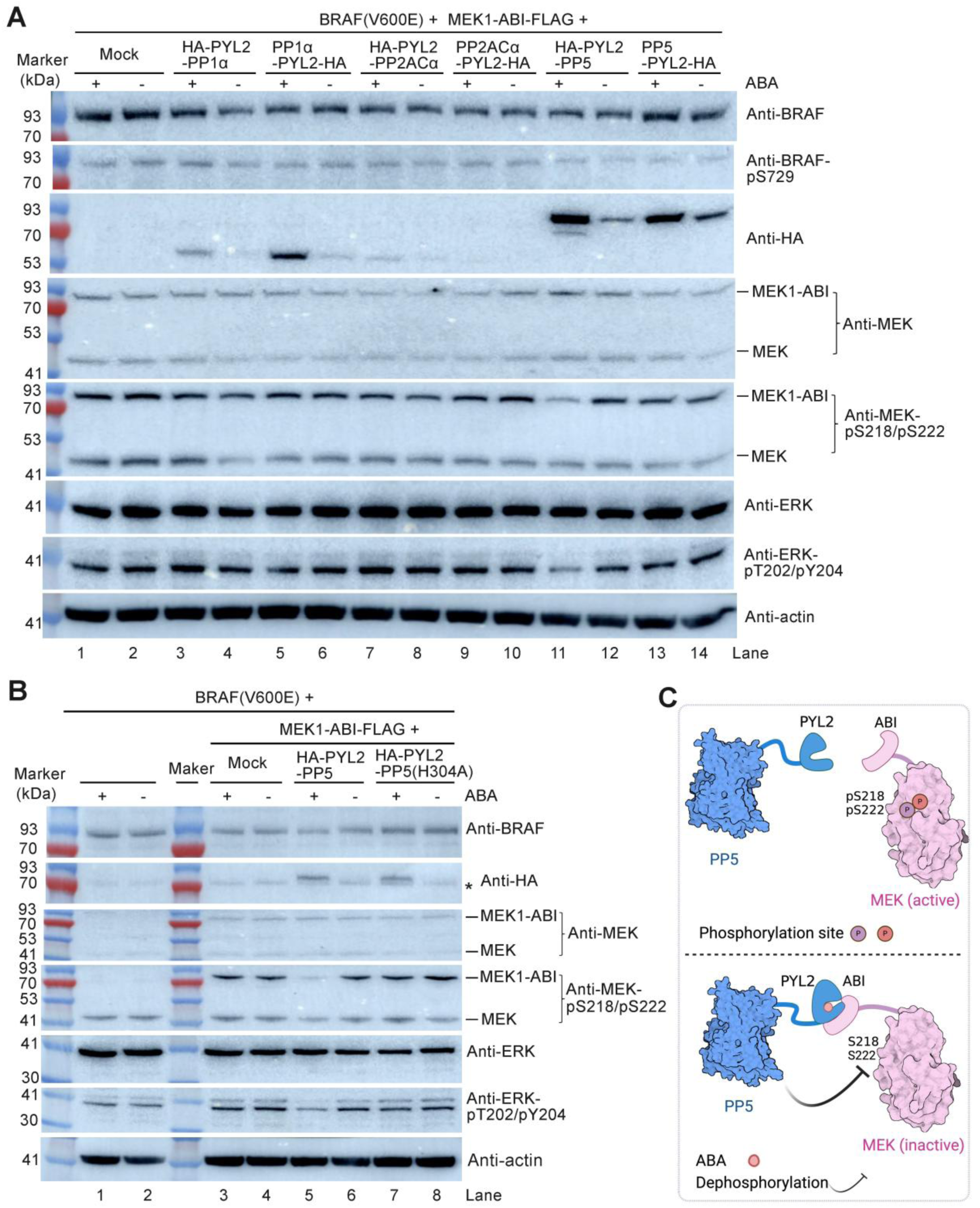
Regulation of MEK1 activity by recruiting protein serine/threonine phosphatases. (A) Effects of PP1α, PP2ACα and PP5 on MEK1 activity when they were recruited to MEK1 in Ba/F3 cells expressing BRAF(V600E) using the ABA induced proximity system. (B) Recruitment of wild-type PP5 but not the catalytic-dead mutant PP5(H304A) to MEK1 in Ba/F3 cells expressing BRAF(V600E) decreased MEK1 at S218/S222 and inhibited the activation of ERK. The asterisk (*) indicates a non-specific band detected by the anti-HA antibody. (C) Illustration of the regulation of MEK by PP5 using the ABA induced proximity system.

### Recruitment of PP5 to BRAF(V600E) cannot facilitate MEK1 dephosphorylation

Since MEK1 can form complex with BRAF^53,54^, we wonder whether recruitment of PP5 to BRAF(V600E) can directly dephosphorylate MEK1. To answer this question, we used purified proteins to reconstitute an *in vitro* assay (Figure 7A). Firstly, BRAF(V600E) with ABI fused to its C terminus was mixed with MEK1 in a kinase reaction buffer at 30 °C for 30 minutes to enable the phosphorylation of MEK1 at S218/S222 (Figure 7B, lanes 1, 2, and 8). Secondly, PP5-PYL2-FLAG in buffers containing palmitoyl-CoA and MnCl_2_ was added and incubated at 30 °C for 50 minutes to catalyze the dephosphorylation of BRAF and MEK1 (Figure 7B, lanes 3 and 9). PP5 alone is in an auto-inhibited state and palmitoyl-CoA was used as an activator of PP5 (ref. ^55^). MnCl_2_ was added because the catalytic activity of PP5 requires the presence of two metal ions, usually Mn^2+^, at its catalytic center^56,57^. In previous studies, 500 µM or even higher concentrations of MnCl_2_ was used to support the activity of purified PP5 in *in vitro* assays^55,57^. However, the cellular concentration of free Mn^2+^ ions is at a low micromolar level^58^. Therefore, we tested two concentrations of MnCl_2_ in the reaction: 50 µM and 500 µM. Thirdly, the reactions were treated with ABA, dabrafenib (DAB, an inhibitor of BRAF), ABA together with DAB, or the same volume of DMSO and incubated at 30 °C for additional 50 minutes (Figure 7B, lanes 4-7 and 10-13). The assay was repeated five times and the phosphorylation of BRAF(V600E) at S729 and that of MEK1 at S218/S222 were quantified by western blotting and normalized to the level of BRAF(V600E) or MEK1 (Figures 7C and 7D).

**Figure 7.**
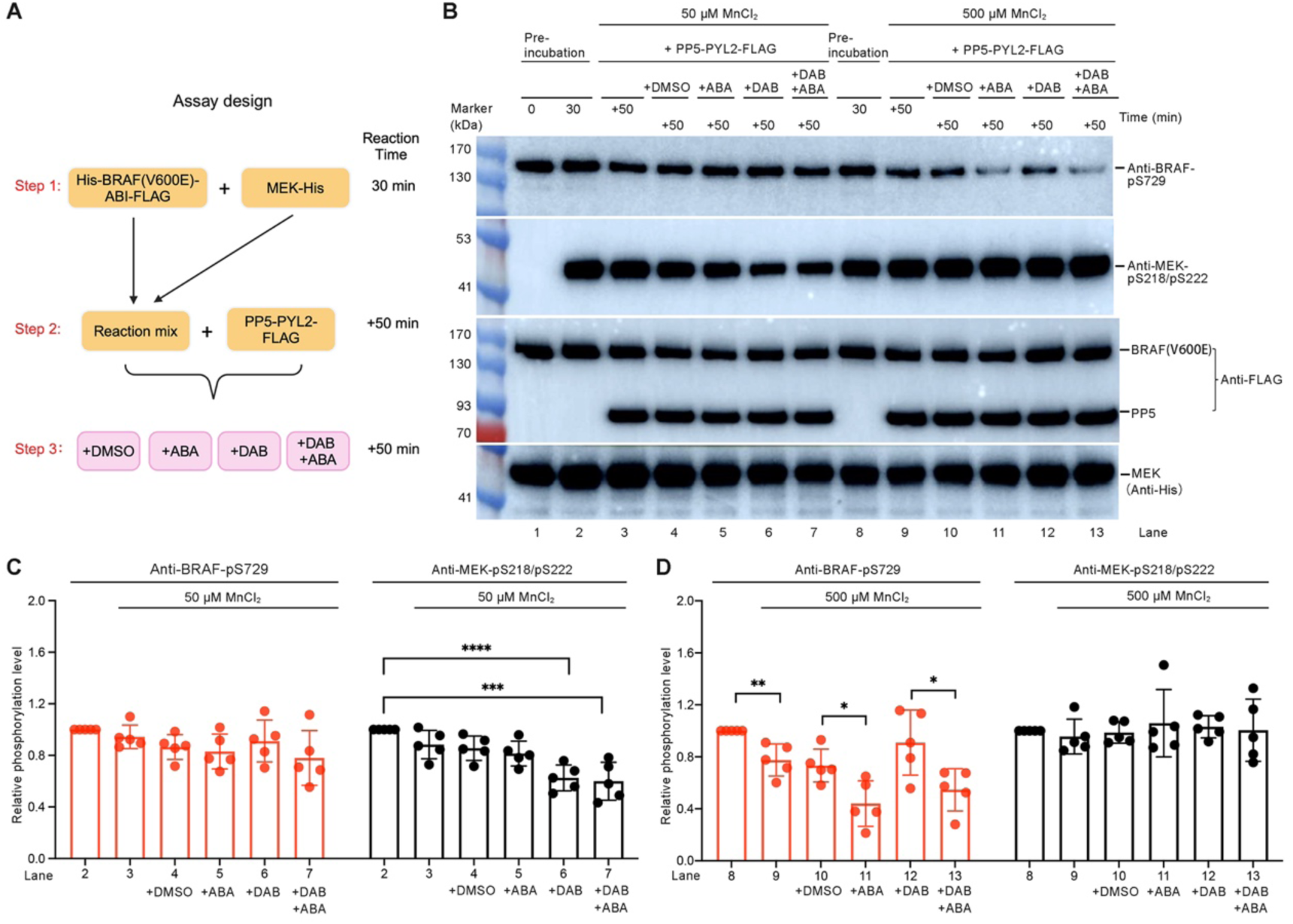
Dephosphorylation of BRAF(V600E) and MEK1 by PP5 can be regulated by Mn^2+^ concentration. (A) Schematic of the assay design. In step 1, purified BRAF(V600E)-ABI with His tag at its N terminus and FLAG tag at its C terminus was first incubated with purified His-tagged MEK1 in a kinase reaction buffer at 30 °C for 30 minutes; in step 2, purified FLAG-tagged PP5-PYL2 was added and incubated for additional 50 minutes in the presence of 50 or 500 µM MnCl_2_; in the step 3, ABA, the BRAF inhibitor Dabrafenib (DAB), ABA plus DAB, or the same volume of DMSO was added and incubated for additional 50 minutes. (B) Analysis of the effects of Mn^2+^ concentration, ABA, and DAB on the activity of PP5 to dephosphorylate BRAF and MEK1 by using western blot. (C and D) Phosphorylation of BRAF(V600E) at S729 and that of MEK1 at S218/S222 were quantified and normalized with their protein bands. Then the normalized phosphorylation signals were further normalized with the phosphorylation signals in lane 2 (50 µM MnCl_2_) or normalized with that in lane 8 (500 µM MnCl_2_). The data represent the mean ± SD of 5 independent measurements and were analyzed using the unpaired *t*-test in Prism to calculate the two-tailed *P* values: \*\*\*\**P* < 0.0001; \*\*\**P* < 0.001; \*\**P* < 0.01; \**P* < 0.05.

In the presence of 50 µM MnCl_2_, the phosphorylation of BRAF(V600E) at S729 was slightly decreased upon adding PP5 but the decrease was not statistically significant regardless of whether ABA or Dabrafenib was added or not (Figure 7C, left panel); the phosphorylation of MEK1 at S218/S222 was also slightly decreased upon adding PP5 and the decrease was also not statistically significant, but upon treatment with Dabrafenib, the phosphorylation of MEK1 at S218/S222 was decreased by 40% (Figure 7C, right panel). In contrast, in the presence of 500 µM MnCl_2_, the phosphorylation of BRAF(V600E) at S729 was significantly decreased after adding PP5 and further decreased upon treatment with ABA (Figure 7D, left panel); the phosphorylation of MEK1 at S218/S222 was not changed after adding PP5 (Figure 7D, right panel).

These results indicate that the recruitment of PP5 to BRAF(V600E) promoted the dephosphorylation of BRAF(V600E), but did not facilitate MEK1 dephosphorylation. Additionally, the phosphatase activity of PP5 is controlled by MnCl_2_ concentration. PP5 dephosphorylated MEK1 at low MnCl_2_ concentration (50 µM), while at high MnCl_2_ concentration (500 µM), PP5 recognized BRAF(V600E) but not MEK1 as its substrate. However, the implication of this finding for the function of PP5 in a cellular context has yet to be studied.

## Discussion

Using the tyrosine kinase BCR::ABL1 and the serine/threonine kinases BRAF(V600E) and MEK1 as examples, we have shown that the ABA induced proximity system provided a convenient method to reversibly control kinase activities and to identify phosphatases that can be recruited to inhibit the kinases. When fusing the ABI and PYL tags to kinases and phosphatases, the positions to which the tags are fused, as well as the length and flexibility of the linkers between ABI and the kinases, and between PYL and the phosphatases, should be taken into consideration. This can help minimize the effect of the tags on the enzymatic activities of the kinases and phosphatases, and can also induce the optimal proximity between the phosphorylation sites of the kinases and related phosphatases, promoting efficient dephosphorylation reactions.

Though the ABA induced proximity system has been studied extensively, the effect of ABA on the stability of PYL2 has not been reported. We have shown that binding of ABA to PYL2 significantly increased the cellular concentration of PYL2. Additionally, when PYL2 was fused to phosphatases such as SHP1 and PP1α, the cellular concentrations of the fusion proteins were also increased significantly upon adding ABA. This unexpected finding suggests that ABA not only induces the proximity of kinases and phosphatases, but also increases the concentrations of phosphatases. Accordingly, we propose that the PYL2-ABA pair can be used to regulate protein concentration in mammalian cells. Since we demonstrated that the protein levels of both FLAG-ABI-BCR::ABL1 and HA-PYL2-SHP1 in the Ba/F3 cells were decreased upon MG132 treatment, and ABA treatment partially restored the protein levels (Figure S3E), it would be interesting to figure out how the protein levels of FLAG-ABI-BCR::ABL1 and HA-PYL2-SHP1 are regulated by MG132 and ABA.

Using the ABA-induced proximity system, we found that BCR::ABL1, the well-known drug target for CML therapy, can be inhibited by the recruitment of several cytoplasmic phosphatases. Additionally, we showed that ABA exhibits a synergistic effect in combination with the BCR::ABL1 inhibitor IMA, emphasizing the importance of developing bifunctional molecules capable of recruiting endogenous cytoplasmic phosphatases to BCR::ABL1 for the treatment of CML.

Another important finding is that the oncogenic kinase BRAF(V600E) and its drug-resistant splicing variants can still be controlled by phosphorylation and dephosphorylation through ABA-induced recruitment of phosphatases. Previous studies have suggested that the oncogenic mutation V600E enables BRAF to bypass the phosphorylation regulation^41–45^. Though phosphorylation can still affect the kinase activity and heterodimerization of BRAF(V600E) with CRAF, it has been reported that phosphorylation has little effect on the transforming activity of BRAF(V600E)^43,59^. In addition, alanine mutations in the phosphorylation sites on the activation loop showed minimal impact on the signaling potential of the BRAF V600F and V600E mutants^45,60^. But we showed that the BRAF(V600E) mediated ERK phosphorylation was downregulated by PP5 but slightly upregulated by PP1α. The phosphatase activities of PP1α and PP5 are necessary for their distinct effects on ERK phosphorylation (Figures 4B and 4C). Recruitment of PP5 to BRAF(V600E) dephosphorylated BRAF(V600E) at multiple Ser/Thr sites. Dephosphorylation of BRAF(V600E) at these sites, except S319/S325, was also observed by Oberoi *et al*. when they treated the HSP90– CDC37–BRAF(V600E) complex with PP5 and quantified the phosphorylation using mass spectrometry^61^. Phosphorylation of BRAF at S365 and T401 has been reported to negatively regulate wild-type BRAF, while phosphorylation at S729 positively regulate wild-type BRAF^43,44^. It has also been reported that mutation of S365 to alanine, mimicking the dephosphorylation state of S365, increased homodimerization of BRAF(V600E) to promote downstream signaling^62^. Therefore, both PP1α and PP5 may have positive and negative effects on the activity of BRAF(V600E) when they are recruited to it. In addition, since BRAF has been reported to form complexes with numerous proteins^9,63^, the distinct effects of PP1α and PP5 on BRAF(V600E) may also be attributed to dephosphorylation of proteins that interact with BRAF(V600E). Further studies are required to unveil the links between the phosphatase activities of PP1α and PP5 and the phosphorylation of ERK.

In addition to the observation that recruitment of PP5 to BRAF(V600E) effectively inhibited the proliferation of Ba/F3 cells expressing BRAF(V600E) or the drug-resistant splicing variants of BRAF(V600E), we have shown that recruitment of PP5 to MEK1 dephosphorylated and inhibited MEK1. The kinase activities of MEK kinases (MEK1 and MEK2) are tightly controlled by phosphorylation. Our results suggest that PP5 can be used to regulate MEK phosphorylation. Development of bifunctional molecules to recruit endogenous PP5 to MEK or RAF kinases is a promising strategy for cancer treatment.

## Acknowledgments

We thank Dr. Hongtao Yu for helpful comments. We thank the reviewers for their constructive feedback. We thank the Mass Spectrometry & Metabolomics Core Facility at the Center for Biomedical Research Core Facilities of Westlake University for sample analysis. This work was supported by the “Pioneer” and “Leading Goose” R&D Program of Zhejiang (2023C03109, 2024SSYS0036), and Westlake Education Foundation.

## Author contributions

Q.H. conceived the project. Y.S., and R.Z. purified the proteins. Y.S. designed and performed all the assays. J.H. and S.F. performed the MS experiments and analyzed the data. Y.S and Q.H. analyzed the data and wrote the manuscript.

## Data and materials availability

Requests for resources and reagents should be directed to the lead contact, Qi Hu. All the data are available in the manuscript or the supplementary materials.

## Supplementary Materials

Materials and Methods Figs. S1 to S6 Table S1

