## Supplementary Information for "Reversible control of kinase signaling through chemical-induced dephosphorylation"

**This PDF file includes:**

Materials and Methods
Figures S1 to S6
Table S1

### Materials and Methods

#### Cell culture

The Ba/F3 cells (from Chinese National Cell Bank, Beijing, China) were cultured in RPMI medium (Hyclone, Cat# SH30809.01) supplemented with 10% FBS (Gibco, Cat# 10270-106), 1% penicillin-streptomycin solution (Hyclone, Cat# SV30010) and 10 ng/mL recombinant mouse IL-3 (GenScript, Cat# Z03111). The 293T cells were cultured in high-glucose DMEM (Hyclone, Cat# C11995500BT) supplemented with 10% FBS and 1% penicillin-streptomycin solution. After cryorecovery, the cells were cultured in a humidified incubator with 5% $CO_2$ at 37 °C for at least two generations before they were used in the assays. The 293F cells were cultured in SMM293-TII medium (Sino Biological, Cat# M293TII) supplemented with 1% penicillin-streptomycin solution at 37 °C under 5% $CO_2$ in a Multitron-Pro shaker (Infors, 110 rpm).

#### Genes and cloning

To build the ABA induced proximity system, we used the *Arabidopsis thaliana* ABI1 (accession number in PubMed: NP_194338.1) and PYL2 (accession number in PubMed: NP_180174.1). Specifically, we used a truncated ABI1 containing residues V126 to L422 with the D143A mutation and called it ABI, and used a truncated PYL2 containing residues T11 to D188 and called it PYL2 in this study.

FLAG-tagged ABI was fused to the N or C termini of kinases including BCR::ABL1 (Addgene plasmid #38158), BRAF (accession number in PubMed: NP_004324.2) with the V600E mutation, aberrantly spliced BRAF(V600E), and MEK1 (accession number in PubMed: NP_002746.1) to obtain the following constructs: FLAG-ABI-BCR::ABL1, FLAG-ABI-BARF(V600E), BRAF(V600E)-ABI-FLAG, spliced BRAF(V600E)-ABI-FLAG, and MEK1-ABI-FLAG. BCR::ABL1 alone, and FLAG-ABI were used as controls. In these constructs, there was a GS linker between the FLAG tag (DYKDDDDK) and ABI, and a SGGGGS linker between ABI and the kinases. To generate retroviruses, the coding sequences of these constructs were separately cloned into MSCV-PGK-puroR vector (Addgene plasmid # 68469); the coding sequence of BRAF(V600E) was cloned into a MSCV-IRES-EGFP vector (Addgene plasmid # 20672).

HA-tagged PYL2 was fused to the N or C termini of phosphatases including SHP1 (accession number in PubMed: NP_002822.2), PTPN5 (accession number in PubMed: NP_008837.1), PTP1B (accession number in PubMed: NP_002818.1), TCPTP (accession number in PubMed: NP_002819.2), PTEN (accession number in PubMed: NP_000305.3), DUSP4 (accession number in PubMed: NP_001385.1), DUSP5 (accession number in PubMed: NP_004410.3), DUSP9 (accession number in PubMed: NP_001305432.1), PP1α (accession number in PubMed: NP_002699.1), PP2ACα (accession number in PubMed: NP_002706.1) and PP5 (accession number in PubMed: NP_006238.1)] to obtain the following constructs: HA-PLY2, HA-PLY2-SHP1, SHP1-PLY2-HA, HA-PLY2(K64A)-SHP1, HA-PLY2-SHP1(C453S), HA-PLY2-PTPN5, HA-PLY2-PTPN5(C496S), PTPN5-PLY2-HA, HA-PLY2-PTP1B, HA-PLY2-PTP1B(C215S), PTP1B-PLY2-HA, HA-PLY2-TCPTP, HA-PLY2-TCPTP(C216S), TCPTP-PLY2-HA, HA-PLY2-PTEN, HA-PLY2-PTEN(C124S), PTEN-PLY2-HA, HA-PLY2-DUSP4, DUSP4-PLY2-HA, HA-PLY2-DUSP5, DUSP5-PLY2-HA, HA-PLY2-DUSP9, DUSP9-PLY2-HA, HA-PLY2-PP1α, HA-PLY2-PP1α(H248K), PP1α-PLY2-HA, HA-PLY2-PP2ACα, PP2ACα-PLY2-HA, HA-PLY2-PP5, PP5-PLY2-HA, HA-PLY2-PP5(H304A), and PP5(H304A)-PLY2-HA. In these constructs, there was a GSG linker between the HA tag (YPYDVPDYA) and PYL2, and a SGGGGSG linker between PYL2 and the phosphatases. To generate retroviruses, the coding sequences of these constructs were separately cloned into a MSCV-PGK-neoR vector.

To construct plasmids for recombinant protein expression, the coding sequences of FLAG-ABI-BCR::ABL1, BRAF(V600E)-ABI-FLAG with an 8x His-tag at its N terminus, and PP5-PYL2-FLAG were separately cloned into the multiple cloning site of a pcDNA3.1 vector. The coding sequence of PYL2-SHP1 with a 6x His-tag at its N terminus was cloned into the multiple cloning site of a modified pET15b vector. The coding sequence of full-length human MEK1 without stop codon was cloned into a pET21b vector.

**Preparation of mouse stem cell virus (MSCV)**

10 mL of the 293T cells (3x $10^5$ cells/mL) were plated in a 10-cm dish. Once the cells were grown to 60% confluency, they were transfected with 7 µg of pCL-Eco (Addgene plasmid # 12371) and 13 µg of MSCV plasmids using 40 µg of Linear Polyethylenimine 25,000 (Polysciences, Cat# 23966) following the manufacturer's instructions (Takara Bio). The medium was exchanged 8 h after transfection. The cells were cultured for 48 h after transfection, then the supernatant was harvested

and filtered through a 0.22 µm syringe filter. The filtrate containing the MSCV retrovirus was used immediately or stored at -80 °C.

#### Generation of stably transduced Ba/F3 cells

To generate Ba/F3 cells stably transduced with MSCV retrovirus carrying the coding sequence of FLAG-ABI, FLAG-ABI-BCR::ABL1, BCR::ABL1, FLAG-ABI-BARF(V600E), BRAF(V600E)-ABI-FLAG, or spliced BRAF(V600E)-ABI-FLAG, 1x $10^6$ Ba/F3 cells per well in 1 mL of RMPI medium containing 20% FBS, 20 ng of mouse IL-3 and 10 µg of polybrene (Merck, Cat# TR-1003-G) were plated into 6-well plates (Corning, Cat# 3516), then 1 mL per well of the retrovirus was added. The Ba/F3 cells were spinfected by centrifugation at 1,000 g for 90 min at 32 °C following the manufacturer's instructions (Takara Bio) and then cultured overnight, followed by replacing the medium with fresh RPMI complete medium (RPMI medium, 10% FBS, 1% P/S) supplied with 10 ng /mL mouse IL-3. 48 h after transduction, the medium was replaced by the RPMI selection medium (RPMI medium, 10% FBS, 1% P/S and 2 µg/mL puromycin) with 10 ng/mL mouse IL-3. The cells were maintained under puromycin (invivogen, Cat# ant-pr) selection for 7 days, passaging as required. Then the transduced cells, except that transduced with FLAG-ABI retrovirus, were cultured in RPMI complete medium without IL-3 to make the cells dependent on the kinase activity of BCR::ABL1 or BRAF(V600E).

To generate Ba/F3 cells stably transduced with BRAF(V600E) retrovirus, 1x $10^6$ Ba/F3 cells per well in 1 mL of RMPI medium containing 20% FBS, 20 ng of mouse IL-3 and 10 µg of polybrene were plated into 6-well plates, then 1 mL per well of the retrovirus was added. The Ba/F3 cells were spinfected by centrifugation at 1,000 g for 90 min at 32 °C following the manufacturer's instructions (Takara Bio) and then cultured overnight, followed by replacing the medium with fresh RPMI complete medium (RPMI medium, 10% FBS, 1% P/S) supplied with 10 ng /mL mouse IL-3. 48 h after transduction, the cells were sorted using FACS (MA900, Sony) based on the EGFP expression. The cells were cultured in RPMI complete medium supplemented with IL-3 for 5 days. The cells were then spun down, washed once with DPBS and resuspended in RPMI complete medium without IL-3 to make the cells dependent on the kinase activity of BRAF(V600E). The stably transduced cells were further transduced with MEK1-ABI-FLAG retrovirus following the same transduction protocol, except that the stably transduced cells were selected using puromycin instead of FACS, and cultured in the absence of IL-3.

The Ba/F3 cells stably transduced with retroviruses carrying FLAG-ABI or kinase coding sequences were further transduced with retroviruses carrying phosphatase coding sequences following similar transduction protocols described above, and the stably transduced cells were selected by treatment with 800 μg/mL of G418 (invivogen, Cat# ant-gn-5) for 7 days. As for the mock groups, the retroviruses carrying empty MSCV vectors were used to transduce the Ba/F3 cells.

**Immunoblotting assay**

ABA (Cat# 862169-250MG) was prepared as a 250 mM stock in DMSO (Sigma, Cat# D2650-100mL). Imatinib (IMA) (Cat# T6230), MG132 (Cat# S2619), and vemurafenib (VEM) (Cat# T8654) were prepared as 100 mM stocks in DMSO. For stably transduced Ba/F3 cells, the cells (2x $10^6$ cells/mL) were transferred to 6-well plates (1 mL/well), followed by adding 2 mL of RPMI complete medium per well supplemented with ABA, IMA, VEM, MG132, or DMSO. Unless otherwise noted, the final concentrations of ABA, IMA and MG132 were 250 μM, 1 μM, and 5 μM, respectively.

The cells were cultured for 12 h, then pelleted by centrifugation (1,000 g, 2 min). The pellet was washed with ice-cold PBS (1 mL x 2), then lysed in RIPA buffer supplemented with protease inhibitors and phosphatase inhibitors on ice for 20 min. The lysates were centrifuged at 22,000 g for 10 min, then the protein concentration in the supernatant was determined by BCA protein assay (Thermo Fisher Scientific, Cat# 23225 and Cat# 23227). The supernatants containing equal amount of total protein (20-40 μg) were mixed with 2x SDS loading buffer, heated at 95 °C for 5 min, subjected to 4-20% SDS-PAGE (GenScript, Cat# M00657) and transferred to 0.2 μm pore size PVDF membrane (Merck Millipore). The membrane was blocked with 5% non-fat milk (Beyotime, Cat# P0216-1500g) in TBST for 1 h at room temperature with shaking, then cut horizontally according to the molecular weight of target proteins, followed by incubation with primary antibodies for 1 h at room temperature or overnight at 4 °C with shaking. The membrane was then washed with TBST for 5 times, incubated with appropriate HRP-conjugated secondary antibodies (Merck Millipore, Cat# AP127P or Cat# AP156P) for 1 h at room temperature, and washed 5 times with TBST. The secondary antibodies on the PVDF membrane were visualized by adding ECL western blot reagents (FDbio, Cat# FD8020), followed by detection of the luminance signal using

Amersham Imager 680.

Primary antibodies used in this study include Anti-actin (Immunoway, Cat# YM3028), Anti-FLAG (Sigma, Cat# F1804), Anti-HA (Abcam, Cat# ab9110), Anti-His (GenScript, Cat# A00186-100), Anti-c-ABL (Thermo Invitrogen™, Cat# MA5-14398), Anti-SHP1 (Abcam, Cat# ab131537), Anti-phospho-Tyr (CST, Cat# 9416S), Anti-STAT5 (CST, Cat# 94205S), Anti-phospho-STAT5 (Tyr694) (CST, Cat# 9359S), Anti-CrkL (Abcam, Cat# ab151791), Anti-phospho-CrkL (Tyr207) (CST, Cat# 3181S), Anti-AKT (CST, Cat# 4691S), Anti-phospho-AKT (Ser473) (CST, Cat# 4060T), Anti-MEK1/2 (CST, Cat# 4694S), Anti-phospho-MEK1/2 (Ser217/221) (CST, Cat# 9154S), Anti-BRAF (CST, Cat# 14814S), Anti-phospho-BRAF (Ser445) (CST, Cat# 2696S), Anti-phospho-BRAF (Ser729) (Abcam, Cat# ab124794), Anti-phospho-BRAF (Thr401) (Abcam, Cat# ab68215), Anti-MAPK (ERK1/2) (CST, Cat# 4695S), Anti-phospho-MAPK (ERK1/2) (Thr202/Tyr204) (CST, Cat# 4370S). The Anti-phospho-MAPK (ERK1/2) (Thr202/Tyr204) antibody detects dually phosphorylated ERK1 (at Thr202 and Tyr204) and ERK2 (at Thr185 and Tyr187), and mono-phosphorylated ERK1 (at Thr202) and ERK2 (at Thr185).

**Cell viability assay**

80 μL per well of the transduced Ba/F3 cells (1x $10^4$ cells) were seeded into 96-well white wall / clear bottom plates (Corning, Cat# 3610) and cultured overnight. Then 20 μL per well of serial dilutions of kinase inhibitors, ABA in RPMI complete medium was added. After additional 48 h, the cell viability was measured by using the CellTiter-Glo® luminescence-based assay (Promega, Cat# G7571). The viability was normalized to that in the DMSO control group. The data from 2 or 3 independent measurements with technical duplicates were analyzed in GraphPad Prism 7. To get the $IC_{50}$ values, the data was fitted using the built-in nonlinear regression method "[Inhibitor] vs. response -- Variable slope (four parameters)".

**Protein expression and purification**

The FLAG-ABI-BCR::ABL1, His-BRAF(V600E)-ABI-FLAG and PP5-PYL2-FLAG were transiently expressed in 293F cells and purified by Anti-FLAG Affinity Resin (GenScript, Cat# L00432-25). 1 L of 293F cells at a density of 2x $10^6$ cells/mL in SMM293-TII medium were transfected with 1-1.5 mg of the pcDNA3.1 plasmid and Linear Polyethylenimine 25,000 (mass

DNA:mass PEI = 1:2). The transfected cells were cultured for 48 h, then harvested by centrifugation (Thermo Fisher SORVALL LYNX 6000 Centrifuge) at 4,000 g for 10 min at 4˚C. The cell pellets were resuspended in the lysis buffer (25 mM Tris 8.0, 150 mM NaCl, 5% glycerol) supplemented with 1.3 μg/mL Aprotinin (VWR, Cat# E429), 1 μg/mL Pepstain (VWR, Cat# J583), 5 μg/mL Leupeptin (VWR, Cat# J580) and 1 mM PMSF (Macklin, Cat# P6140), and lysed by ultrasonication (SONICS vibra cell$^{TM}$). The cell lysates were centrifuged at 22,000 g at 4 °C for 1 h (BECKMAN Avanti JXN-26 Centrifuge), then the target proteins in the supernatants were purified by using anti-FLAG affinity resin following the manufacturer's protocol. For FLAG-ABI-BCR::ABL1, it was further purified by a Source-15Q (GE Healthcare). The eluates from anti-FLAG affinity resin or Source-15Q column were supplemented with 5 mM Dithiothreitol (DTT) and further purified by Superdex 200 increase 10/300 GL column (GE Healthcare) and eluted by SEC buffer (25 mM HEPES 7.4, 150 mM NaCl). The peak fractions were pooled and concentrated for biochemical assays.

The His-PYL2-SHP1 and MEK-His were overexpressed in *Escherichia coli* BL21(DE3) and purified by TALON Metal Affinity Resin (TaKaRa, Cat# 635504). The plasmids were transformed into BL21(DE3) cells. The cells were cultured in LB medium supplemented with 0.1 mg/mL ampicillin at 37 °C until OD600 reached 1-1.5, then cooled to 20 °C followed by addition of 200 μM β-D-thiogalactopyranoside (IPTG). Then the cells were cultured at 20 °C overnight. The cells were harvested by centrifugation at 4,000 g for 10 min at 4 °C and resuspended in the lysis buffer (25 mM Tris 8.0, 150 mM NaCl, 5% glycerol) supplemented with 1 mM PMSF and lysed by ultrasonication. The cell lysates were centrifuged at 22,000 g at 4 °C for 1 h (BECKMAN Avanti JXN-26 Centrifuge), then the target proteins in the supernatants were purified by using TALON Metal Affinity Resin following the manufacturer's protocol. The eluates were supplemented with 5 mM Dithiothreitol (DTT) and loaded into a Source-15Q column (GE Healthcare) and eluted by a linear gradient from 100% buffer A (25 mM Tris 8.0, 5% glycerol) to 40% Buffer B (25 mM Tris 8.0, 1 M NaCl, 5% glycerol). The peak fractions were pooled, supplemented with 5 mM DTT, and further purified by Superdex 200 increase 10/300 GL column (GE Healthcare) and eluted by SEC buffer (25 mM HEPES 7.4, 150 mM NaCl). The peak fractions were pooled and concentrated for biochemical assays.

**Gel filtration assay**

Gel filtration assay was used to analyze the binding of PYL2-SHP1 to ABI-BCR::ABL1 in the presence of ABA. 100 μL of purified His-PYL2-SHP1 (10 μM) in SEC buffer (25 mM HEPES 7.4, 150 mM NaCl) was mixed with 100 μL of purified FLAG-ABI-BCR::ABL1 (1 μM) in SEC buffer, or with 100 μL of SEC buffer alone. The resulting 200 μL mixture was incubated with 100 μL of ABA (750 μM), which was prepared by diluting a 250 mM stock in DMSO using SEC buffer, or mixed with 100 μL of a DMSO control in SEC buffer at 4 °C for 4 hours. Each of these mixtures was subsequently subjected to gel filtration using the Superdex 200 increase 10/300 GL column (GE Healthcare). The eluted fractions were collected and analyzed by SDS-PAGE.

***In vitro* dephosphorylation assay of BCR::ABL1**

Purified FLAG-ABI-BCR::ABL1 at a concentration of 0.6 μM in 25 mM HEPES pH 7.4, 150 mM NaCl was diluted to 60 nM using buffer K (50 mM HEPES pH 7.4, 150 mM NaCl, 0.01% Triton X-100, 0.01% BSA, 5 mM $MgCl_2$ and 2 mM DTT). Purified His-PYL2-SHP1 at a concentration of 10 μM in 25 mM HEPES pH 7.4, 150 mM NaCl was diluted to 120 nM using buffer K. ATP at pH 7.0 at a concentration of 100 mM in $H_2O$ was diluted to 400 μM using buffer K. 150 μL of the diluted FLAG-ABI-BCR::ABL1 was mixed with 75 μL of the diluted His-PYL2-SHP1, then 75 μL of the diluted ATP was added to initiate the phosphorylation reaction. The reaction was carried out at 30 °C in a metal bath. At time points of 0 min and 50 min, 12.5 μL of the reaction mixture was transferred to 2x SDS loading buffer and immediately heated at 95 °C for 5 min to stop the reaction.

After 50 min, 100 μL of the reaction mixture was mixed with 100 μL of 500 μM ABA that was prepared by dilution of a 250 mM stock in DMSO using buffer K, or mixed with 100 μL of a DMSO control in buffer K. At time points of additional 20 min and 50 min, 25 μL of the reaction mixture was removed and mixed with 2x SDS loading buffer and immediately heated at 95 °C for 5 min. The samples mixed with SDS loading buffer were then subjected to immunoblotting.

***In vitro* dephosphorylation assay of BRAF(V600E) and MEK**

Purified His-BRAF(V600E)-ABI-FLAG at a concentration of 5 μM in 25 mM HEPES 7.4, 150 mM NaCl was diluted to 240 nM using buffer A (50 mM HEPES 7.4, 150 mM NaCl, 0.01% Triton X-100, 0.01% BSA, 2 mM $MgCl_2$, 2 mM DTT). MEK-His at a concentration of 56 μM in 25 mM

HEPES 7.4, 150 mM NaCl was diluted to 1.2 μM using buffer A. ATP at pH 7.0 at a concentration of 100 mM in $H_2O$ was diluted to 400 μM using buffer A. PP5-PYL2-FLAG at a concentration of 7 μM in 25 mM HEPES 7.4, 150 mM NaCl was diluted to 150 nM in buffer B (50 mM HEPES 7.4, 150 mM NaCl, 0.01% Triton X-100, 0.01% BSA, 100 μM $MnCl_2$, 2 mM DTT, 100 μM Palmitoyl-CoA) or buffer C (50 mM HEPES 7.4, 150 mM NaCl, 0.01% Triton X-100, 0.01% BSA, 1 mM $MnCl_2$, 2 mM DTT, 100 μM Palmitoyl-CoA). Palmitoyl-CoA was purchased from Sigma (Cat# P9716) and prepared as a 10 mM stock in $H_2O$. Dabrafenib (TargetMol, Cat# T1903) was prepared as a 10 mM stock in DMSO.

Firstly, 160 μL of the diluted His-BRAF(V600E)-ABI-FLAG was mixed with 160 μL of the diluted MEK-His. The kinase reaction was initiated by adding 320 μL of the diluted ATP and then carried out at 30 °C. At time points of 0 min and 30 min, 5 μL of the reaction mixture was removed and immediately mixed with 2x SDS loading buffer and heated at 95 °C for 5 min. Secondly, 250 μL of the reaction mixture was mixed with 250 μL of the diluted PP5-PYL2-FLAG in buffer B or C to initiate the dephosphorylation reaction. The reaction was carried out at 30 °C for 50 min, then 10 μL of the reaction mixture was removed and mixed with 2x SDS loading buffer and heated at 95 °C for 5 min. Thirdly, 384 μL of the remaining reaction mixture was transferred to four Eppendorf tubes (96 μL per tube); to each tube, 24 μL of ABA (1250 μM), 24 μL of Dabrafenib (50 μM), or 24 μL of ABA (1250 μM) plus Dabrafenib (50 μM), or 24 μL of a DMSO control, in buffer D (50 mM HEPES 7.4, 150 mM NaCl, 0.01% Triton X-100, 0.01% BSA, 1 mM $MgCl_2$, 50 μM $MnCl_2$, 2 mM DTT and 50 μM Palmitoyl-CoA) or in buffer E (50 mM HEPES 7.4, 150 mM NaCl, 0.01% Triton X-100, 0.01% BSA, 1 mM $MgCl_2$, 500 μM $MnCl_2$, 2 mM DTT and 50 μM Palmitoyl-CoA) was added. After additional 50 min, 12.5 μL of each reaction mixture was removed and mixed with 2x SDS loading buffer and heated at 95˚C for 5 min.

The samples mixed with SDS loading buffer were subjected to immunoblotting. The bands on the immunoblotting image were quantified using ImageJ 1.53k.

**Mass spectrometry**

50 mL of the Ba/F3 cells stably expressing FLAG-tagged kinase-ABI or ABI-kinase fusion protein together with HA-tagged PYL2-phosphatase fusion protein, were treated with 50 μL of ABA (250 mM stock in DMSO) or 50 μL of DMSO for 12 h. Then the cells were pelleted by centrifugation

(1,000 g, 2 min), washed twice with ice-cold PBS, lysed by incubating with 800 μL of the lysis buffer (25 mM Tris 8.0, 150 mM NaCl, 5% glycerol) supplemented with 1% Triton X-100, 1 mM EDTA, and protease inhibitors and phosphatase inhibitors at 4 °C for 1.5 h. The lysates were centrifuged at 22,000 g for 10 min. The protein concentration in the supernatant was determined by BCA protein assay. 70 μL of the supernatant was mixed with 2x SDS loading buffer, heated at 95 °C for 5 min and then subjected to immunoblotting.

About 700 μL of the supernatant was incubated with 50 μL of MonoRab™ Anti-DYKDDDDK Magnetic Beads (GeneScript, Cat# L00835-1) at 4 °C for 4 h, washed and eluted following the manufacturer's protocol. The eluate (about 200 μL) was mixed with 50 μL of 5x SDS loading buffer, separated by SDS-PAGE, and stained with Coomassie brilliant blue G-250. The gel bands of interest were cut into pieces. Proteins in the gel were reduced and alkylated in 50 mM ammonium bicarbonate at 37 °C overnight and then digested by trypsin. The digested products were extracted twice with 0.1% formic acid in 50% acetonitrile aqueous solution and dried by SpeedVac.

For LC-MS/MS analysis, the digested products were loaded into an analytical column made by packing C-18 resin (300 A, 3 μm, Varian, Lexington, MA) into a fused silica capillary column (75 μm ID, 150 mm length; Upchurch, Oak Harbor, WA), and then eluted by a linear gradient from 99% mobile phase A (0.1% formic acid in water) to 99% mobile phase B (80% acetonitrile and 0.1% formic acid) over 120 min (for the detection of protein phosphorylation) or 65 min (for the detection of phosphatases that were recruited to BRAF of MEK) with a flow rate of 0.300 μL/min in a Thermo Vanquish Neo integrated nano-HPLC system that was directly interfaced with a Thermo Exploris 480 mass spectrometer. The mass spectrometer was operated in the data-dependent acquisition mode using the Xcalibur 4.1 software. A single full-scan mass spectrum in the Orbitrap (350-1800 m/z, 60,000 resolution) was done followed by several data-dependent MS/MS scans at 30% normalized collision energy. The AGC target was set as 5e4, and the maximum injection time was 50 ms. Each mass spectrum was analyzed using the Thermo Xcalibur Qual Browser and Proteome Discoverer (2.5.0.400). The Sequest search parameters included a 10 ppm precursor mass tolerance, 0.02 Da fragment ion tolerance, and up to 2 internal cleavage sites. Cysteine alkylation was treated as fixed modification. Methionine oxidation and STY phosphorylation were treated as variable modifications. Peptides were filtered with 1% false discovery rate (FDR).

### Supplementary Figures

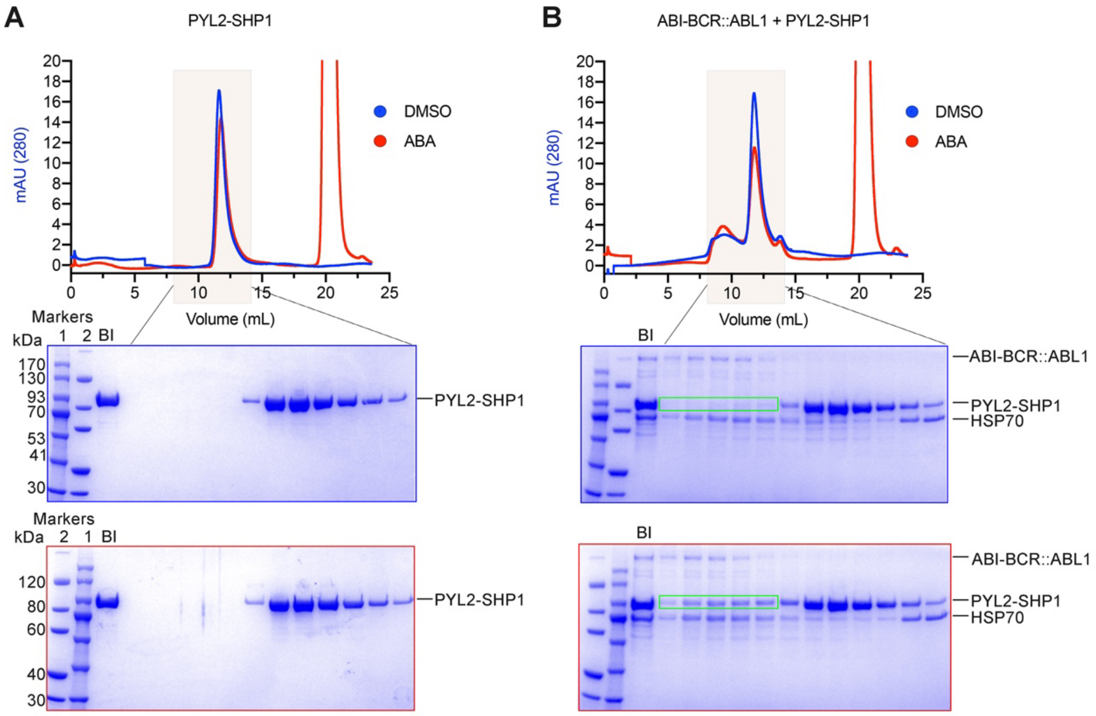


**Figure S1. ABA-induced recruitment of PYL2-SHP1 to ABI-BCR::ABL1.**

**(A)** His-PYL2-SHP1 was first incubated with ABA or DMSO, then separated by gel filtration, analyzed by SDS-PAGE, and stained by Coomassie blue. The gels with blue and red borders correspond to samples incubated with DMSO and ABA, respectively. BI: before injection.

**(B)** His-PYL2-SHP1 mixed with FLAG-ABI-BCRABL was first incubated ABA of DMSO, then analyzed by the gel filtration assay. The green boxes indicate the bands of PYL2-SHP1 that shifted together with ABI-BCR::ABL1 in the presence of ABA but not DMSO.

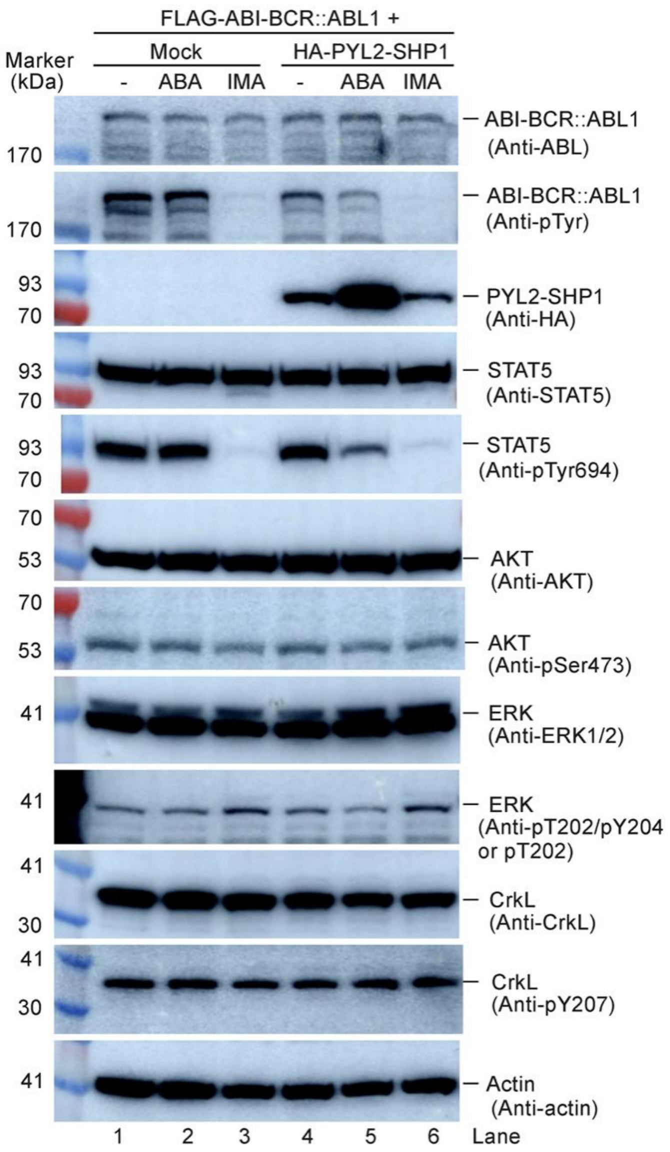


**Figure S2. Effects of ABA-induced recruitment of PYL2-SHP1 to ABI-BCR::ABL1 and inhibition of BCR::ABL1 by imatinib (IMA) on the downstream signaling pathways.**

Both ABA treatment and IMA treatment inhibited the phosphorylation of ABI-BCR::ABL1 and decreased the phosphorylation level of STAT5 downstream of BCR::ABL1, but had a moderate effect on the phosphorylation of ERK and had little effect on the phosphorylation of AKT and CrkL.

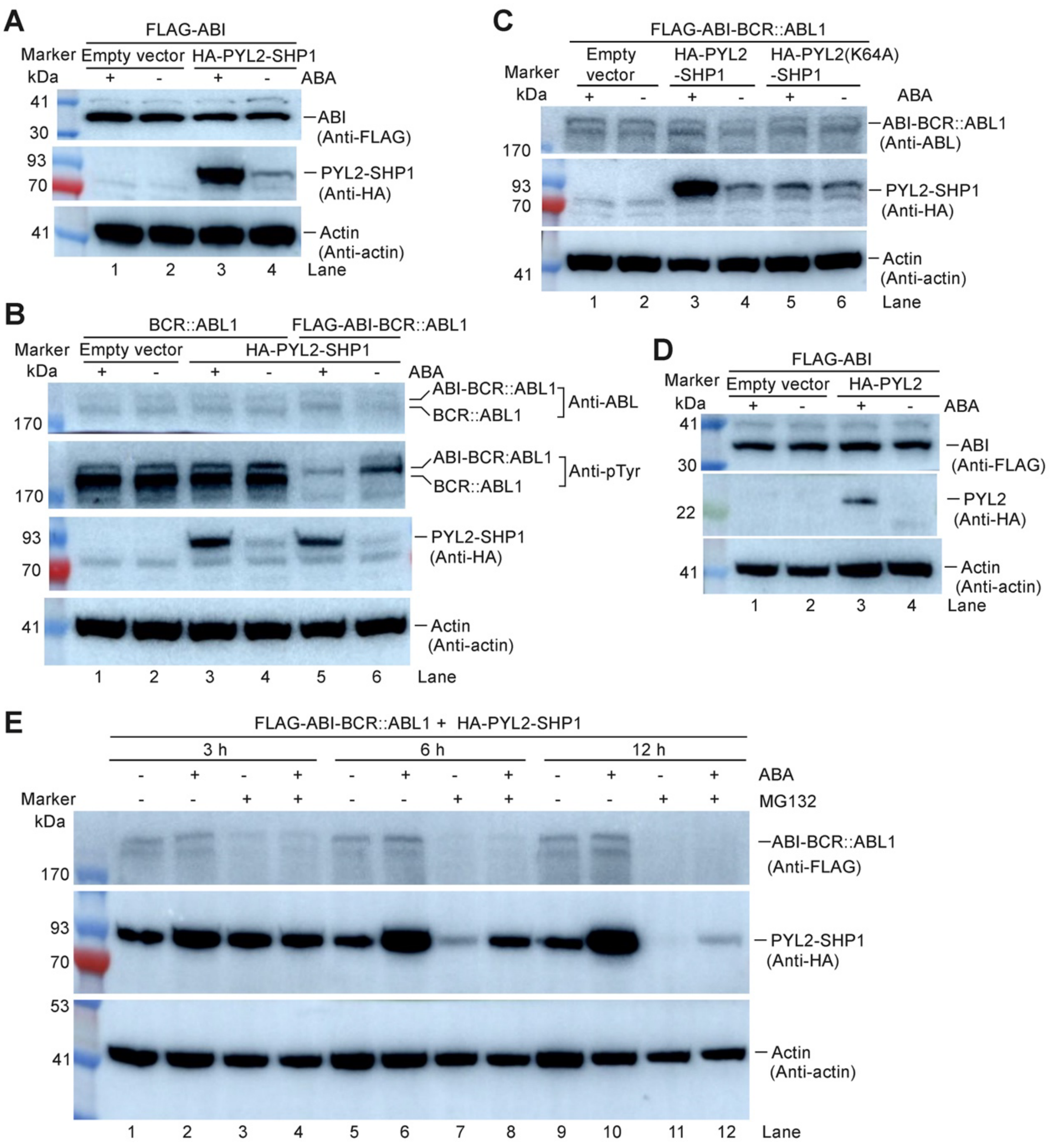


**Figure S3. The binding of ABA to PYL2 upregulated the protein level of PYL2 and its fusion protein with SHP1 in Ba/F3 cells.**

**(A)** The protein level of HA-PYL2-SHP1 was increased upon ABA treatment in Ba/F3 cells stably expressing both FLAG-tagged ABI and HA-PYL2-SHP1.

**(B)** Without recruitment of BCR::ABL1, ABA treatment still upregulated the protein level of HA-PYL2-SHP1 in Ba/F3 cells.

**(C)** With a K64A mutation in PYL2 blocking its binding to ABA, adding ABA could no longer increase the protein level of HA-PYL2-SHP1 in Ba/F3 cells.

**(D)** Without SHP1 fused to PYL2, ABA treatment still increased the protein level of HA-tagged PYL2 in Ba/F3 cells.

(**E**) MG132 treatment decreased the protein levels of both FLAG-ABI-BCR::ABL1 and HA-PYL2-SHP1 in Ba/F3 cells.

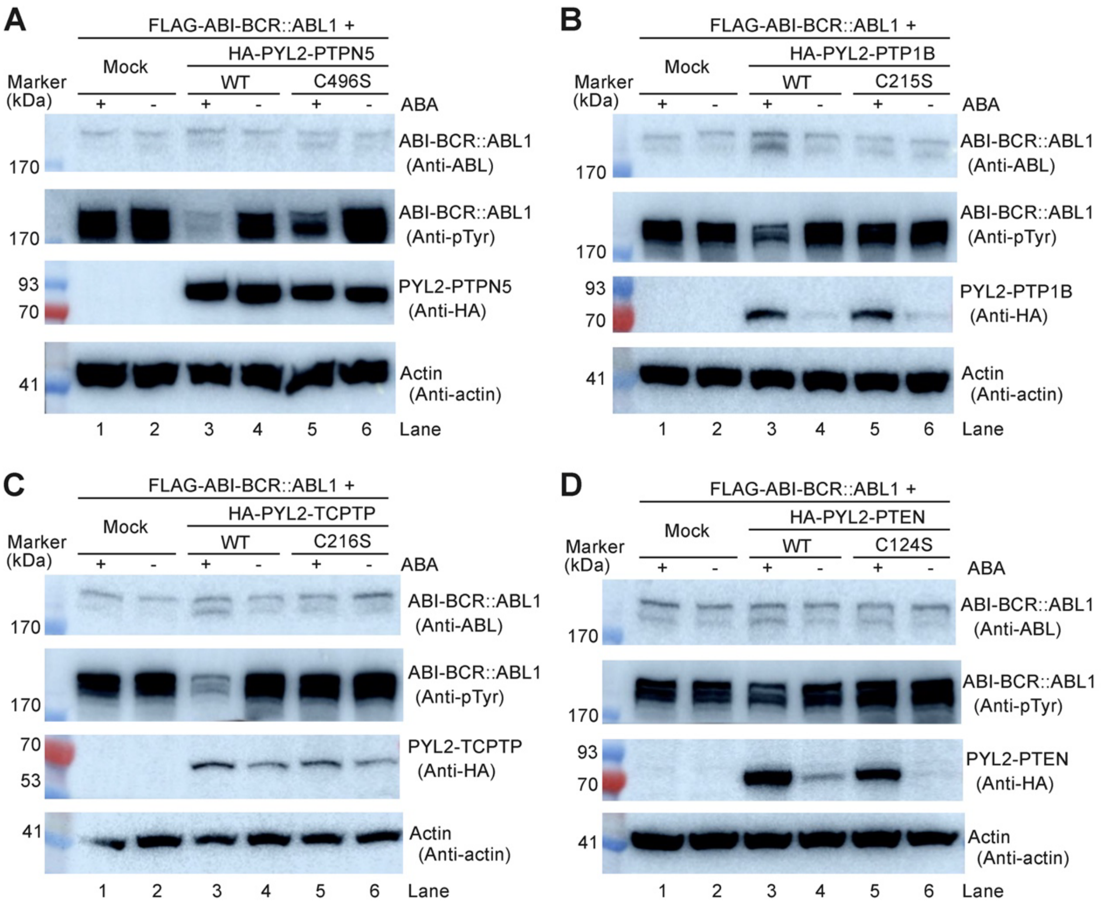


**Figure S4. Dephosphorylation of BCR::ABL1, induced by recruitment of phosphatases, is dependent on the catalytic activities of these phosphatases.**

Recruitment of HA-tagged PYL2-PTPN5 **(A)**, PYL2-PTP1B **(B)**, PYL2-TCPTP **(C)**, or PYL2-PTEN **(D)**, but not the catalytic-dead mutants of these phosphatases, to FLAG-tagged ABI-BCR::ABL1 in Ba/F3 cells significantly decreased the pTyr level of BCR::ABL1.

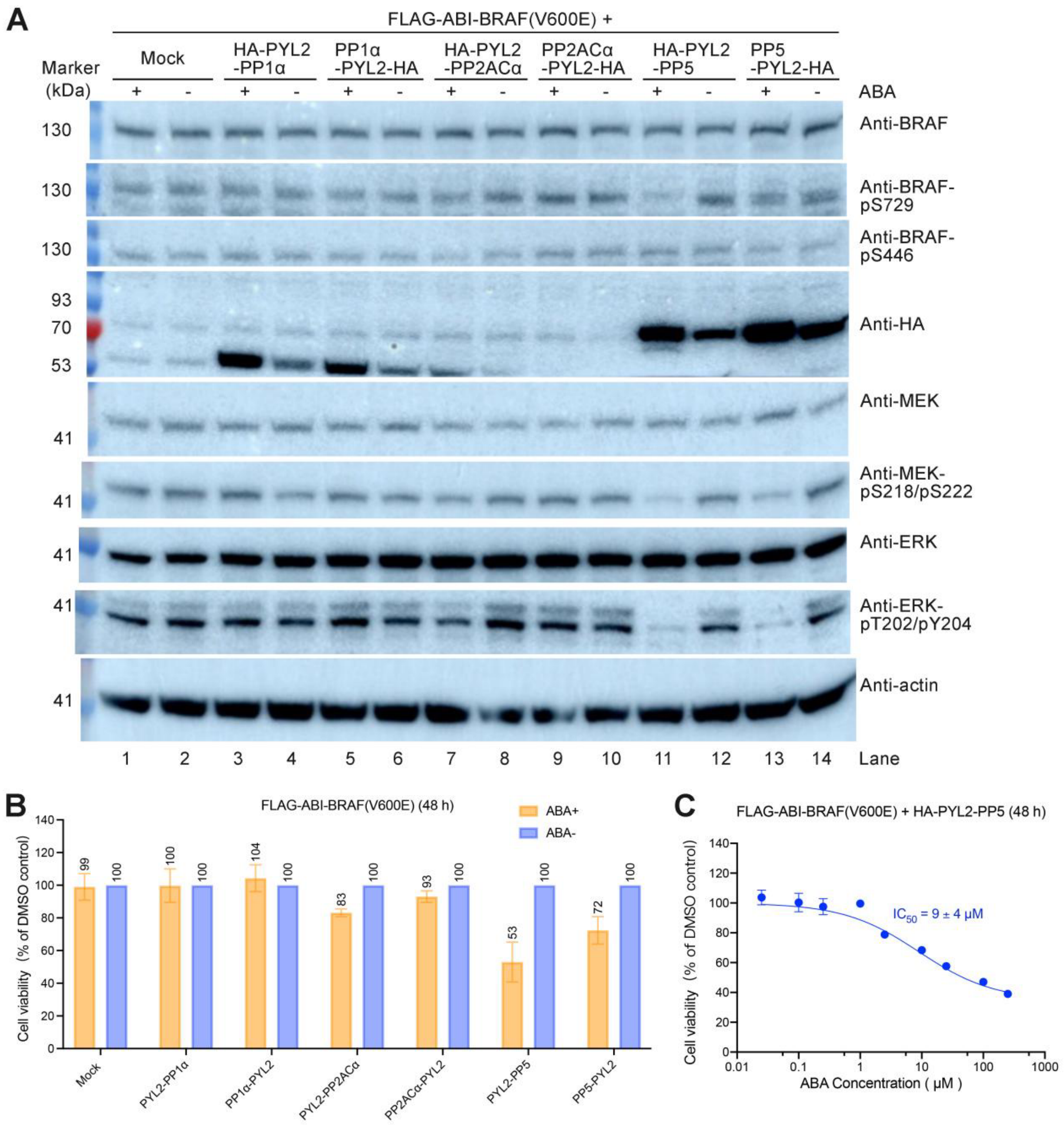


**Figure S5. Regulation of BRAF(V600E) activity by recruiting protein serine/threonine phosphatases.**

**(A)** Recruitment of fusion proteins of PYL2 with PP1α, PP2ACα and PP5 to FLAG-ABI-BRAF(V600E) by adding ABA in Ba/F3 cells showed distinct effects on the BRAF(V600E)-MEK-ERK signaling pathway.

**(B)** Inhibition of the viability of Ba/F3 cells expressing FLAG-ABI-BRAF(V600E) by recruitment of PP1α, PP2ACα or PP5 to BRAF(V600E). The viability of the ABA-treated cells was normalized to that of the untreated cells. The values represent the mean ± SD of three independent measurements with technical duplicates.

**(C)** ABA inhibited the viability of Ba/F3 cells expressing FLAG-ABI-BRAF(V600E) and HA-

PYL2-PP5, with an $IC_{50}$ of 9.0 μM. The values represent the mean ± SD of two independent measurements with technical duplicates.

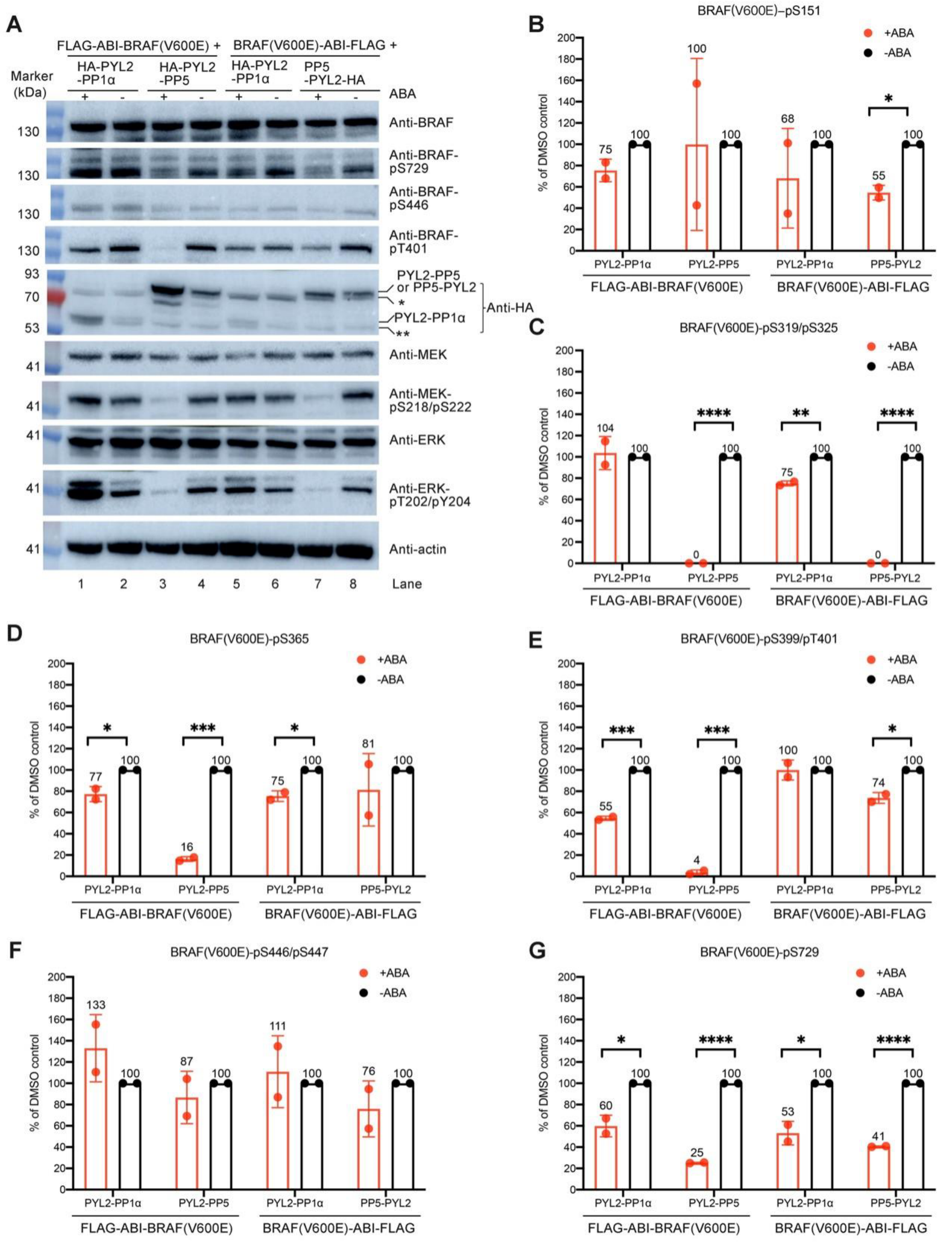


**Figure S6. Analysis of BRAF(V600E) phosphorylation in Ba/F3 cells using mass spectrometry**

**(A)** Recruitment of PP1α and PP5 to BRAF(V600E) using the ABA induced proximity system in Ba/F3 cells showed distinct effects on the BRAF(V600E)-MEK-ERK signaling pathway.

**(B-G)** Quantification of the effect of ABA treatment on the phosphorylation of BRAF(V600E) in Ba/F3 cells stably expressing FLAG-ABI-BRAF(V600E) together with HA-PYL2-PP1α or with HA-PYL2-PP5, or stably expressing BRAF(V600E)-ABI-FLAG together with HA-PYL2-PP1α or with PP5-

PYL2-HA. The abundance of each targeted peptide was normalized to the average of total peptides abundance of BRAF(V600E) in the same group. Then the normalized abundance was normalized to that in the corresponding DMSO control group. The data represent the mean ± SD of two independent measurements and were analyzed using the multiple unpaired *t* tests in Prism to calculate the two-tailed *P* values: ****$P < 0.0001$; ***$P < 0.001$; **$P < 0.01$; *$P < 0.05$.

**Table S1. Identification of PP1 and PP2A subunits recruited to BRAF(V600E) and MEK in the presence of ABA by using mass spectrometry (MS).**

| Sample | PP1 and PP2A subunits identified by MS | Abundance (x $10^6$) | | | | | | | |
|---|---|---|---|---|---|---|---|---|---|
| | | 1st experiment | | 2nd experiment | | 3rd experiment | | 4th experiment | |
| | | ABA | DMSO | ABA | DMSO | ABA | DMSO | ABA | DMSO |
| BRAF(V600E)-ABI-FLAG + HA-PYL2-PP1α | PPP1CA (PP1α) | 114.05 | 9.84 | 295.80 | 74.88 | 353.64 | 168.18 | 403.96 | 181.42 |
| | Ppp1r18 | 81.06 | 54.68 | 33.36 | 5.86 | 32.08 | 8.50 | 16.24 | 0.00 |
| BRAF(V600E)-ABI-FLAG + HA-PYL2-PP2ACα | PPP2CA (PP2ACα) | 52.41 | 21.25 | 86.83 | 40.97 | 51.86 | 32.07 | 84.26 | 38.18 |
| | Ppp2r1b | 4.93 | 1.03 | 7.20 | 0.00 | 6.28 | 1.38 | 5.65 | 1.34 |
| MEK-ABI-FLAG + HA-PYL2-PP1α | PPP1CA (PP1α) | 299.28 | 16.41 | 756.53 | 116.98 | 576.55 | 255.88 | 600.14 | 269.71 |
| | Ppp1r18 | 119.15 | 38.29 | 128.97 | 5.15 | 257.66 | 251.90 | 94.04 | 20.74 |
| | Ppp1r13l | 5.24 | 1.18 | 17.12 | 4.42 | 6.24 | 3.65 | 8.85 | 3.33 |
| | Ppp1r7 | 7.97 | 0.00 | 27.27 | 2.02 | 6.97 | 1.81 | 7.65 | 0.81 |
| MEK-ABI-FLAG + HA-PYL2-PP2ACα | PPP2CA (PP2ACα) | 99.13 | 31.23 | 140.68 | 55.33 | 88.49 | 42.02 | 119.28 | 32.49 |
| | Ppp2r1b | 0.82 | 0.77 | 4.53 | 4.51 | 7.22 | 4.54 | 5.99 | 2.13 |
